# A transcriptional cycling model recapitulates chromatin-dependent features of noisy inducible transcription

**DOI:** 10.1101/2022.05.03.490387

**Authors:** M. Elise Bullock, Nataly Moreno-Martinez, Kathryn Miller-Jensen

## Abstract

Activation of gene expression in response to environmental cues results in substantial phenotypic heterogeneity between cells that can impact a wide range of outcomes including differentiation, viral activation, and drug resistance. An important source of gene expression noise is transcriptional bursting, or the observation that transcripts are produced during infrequent bursts of promoter activity. Chromatin accessibility, which regulates assembly of polymerase complexes on promoters, impacts transcriptional bursting, suggesting that how an activating signal affects transcriptional noise will depend on the initial chromatin state at the promoter. To explore this possibility, we simulated transcriptional activation using a transcriptional cycling model with three promoter states that represent chromatin remodeling, polymerase binding and pause release. We initiated this model over a large parameter range representing target genes with different chromatin environments, and found that, upon increasing the polymerase pause release rate to activate transcription, changes in gene expression noise varied significantly across initial promoter states. This model captured phenotypic differences in activation of latent HIV viruses integrated at different chromatin locations and mediated by the transcription factor NF-κB. Activating transcription in the model via increasing one or more of the transcript production rates, as occurs following NF-κB activation, reproduced experimentally measured transcript distributions for four different latent HIV viruses, as well as the bimodal pattern of HIV protein expression that leads to a subset of reactivated virus. Importantly, the parameter ‘activation path’ differentially affected gene expression noise, and ultimately viral activation, in line with experimental observations. This work demonstrates how upstream signaling pathways can be connected to biological processes that underlie transcriptional bursting, resulting in target gene-specific noise profiles following stimulation of a single upstream pathway.

**Author Summary:** Many genes are transcribed in infrequent bursts of mRNA production through a process called transcriptional bursting, which contributes to variability in responses between cells. Heterogeneity in cell responses can have important biological impacts, such as whether a cell supports viral replication or responds to a drug, and thus there is an effort to describe this process with mathematical models to predict biological outcomes. Previous models described bursting as a transition between an “OFF” state or an “ON” state, an elegant and simple mathematical representation of complex molecular mechanisms, but one which failed to capture how upstream activation signals affected bursting. To address this, we added an additional promoter state to better reflect biological mechanisms underlying bursting. By fitting this model to variable activation of quiescent HIV infections in T cells, we showed that our model more accurately described viral expression variability across cells in response to an upstream stimulus. Our work highlights how mathematical models can be further developed to understand complex biological mechanisms and suggests ways to connect transcriptional bursting to upstream activation pathways.

## Introduction

Heterogeneity in gene expression between cells impacts a wide range of phenotypic outcomes, including differentiation [1, 2], viral expression [3–7], and drug resistance [8]. In eukaryotic cells, a major source of gene expression heterogeneity is transcriptional bursting [9–11], in which a gene promoter infrequently produces bursts of transcripts. Two metrics are often used to describe a gene’s burstiness: burst size, quantifying how many transcripts are produced in one burst, and burst frequency, describing how many bursts occur over time. Influencing transcriptional bursting, either through altering burst size or burst frequency, can aid in clinically relevant settings where transcriptional noise plays a role in disease progression [12].

Several possible points of control have been proposed for transcriptional bursting, including transcription factor regulation [13, 14], polymerase recycling [15], chromatin environment [11, 16], nucleosome positioning [17, 18], and polymerase pause regulation [4, 19]. Despite this complexity, transcriptional bursting is most often modeled as a simple random telegraph process, in which a promoter infrequently transitions from an “OFF” state to an “ON” state [20]. Regardless of this simplicity, the two-state model accurately reflects transcriptional bursting in many biological contexts [11, 21]. To address situations in which the two-state model does not capture all aspects of transcriptional noise, additional layers of complexity have been added, including multiple “OFF” states [20,22,23], a continuum of states [24], and a refractory state [11,25,26].

One biological context in which the two-state promoter model lacks descriptive detail is in recapitulating the role of transcription factor (TFs) activation in inducible gene expression. While several studies have assessed how TFs modulate inducible gene expression noise [27, 28], they are limited by an inability to directly connect molecular steps in transcription to changes in promoter state. TFs often recruit molecular complexes that alter aspects of both transcriptional burst size and burst frequency. For example, tumor necrosis factor (TNF) initiates a signaling cascade that activates the canonical TF nuclear factor-κB (NF-κB) [29]. NF-κB mediates recruitment of histone acetyltransferases (HATs), including CBP/p300 [30], to target promoters, destabilizing DNA-histone interactions within nucleosomes and increasing promoter accessibility. NF-κB also mediates recruitment of Mediator, RNA polymerase II (RNAPII) and other members of the preinitiation complex to accessible promoters [31]. Finally, NF-κB interacts with bromodomain-containing protein 4 (Brd4) to recruit and activate the positive elongation factor b (P-TEFb), releasing paused RNAPII and allowing efficient transcriptional elongation to proceed [32–34]. These processes are important for inducible transcription in many contexts including inflammatory gene expression [35] and latent HIV activation [4, 36].

A recent model of transcriptional bursting that includes an additional promoter state to decouple RNAPII pausing from RNAPII recruitment more accurately describes steady-state transcriptional bursting [7, 19]. Here, we explored if this three-state promoter model of transcription, which explicitly models the transcriptional steps of chromatin remodeling, polymerase recruitment, and polymerase pause release, could also capture features of noisy inducible transcription. Through deterministic and stochastic simulations, we explored parameter ranges for a three-state promoter model that describe chromatin environments of quiescent-but-inducible promoters in a range of biological contexts. We found that upon transcriptional activation implemented by increasing the polymerase pause release rate, changes in gene expression noise varied significantly across initial promoter states. We then fit our model simulations to time-resolved, single-cell experimental data of how NF-κB activation induces transcription of latent HIV viruses integrated into different chromatin environments. We found that the model accurately captured how exogenous stimulation of NF-κB differentially affected transcriptional noise and viral reactivation initiated in a range of basal states. Furthermore, the model allowed for exploration of the influence of multiple NF-κB-mediated steps in transcriptional activation. We anticipate this model will offer a means to connect transcriptional bursting to upstream signaling pathways, with applications that extend to other NF-κB-mediated biological systems, and potentially other signaling pathways.

## Results

### Steady-state analysis of a transcriptional cycling model reveals that the BIR:BTR ratio controls promoter remodeling and the PPRR:PBR ratio controls transcriptional cycling

To explicitly consider the effect of promoter accessibility and RNAPII pausing on transcriptional noise, we explored a previously published three-state promoter model [19]. In this model, transcription of a target gene is regulated by its promoter which transitions between three states, only one of which is productive for producing mRNA (Fig 1A). The rate at which the unavailable promoter (UP) state transitions to the available promoter (AP) state is the burst initiation rate (BIR) and the returning promoter transition rate is the burst termination rate (BTR). These two rates could be assumed to describe the remodeling of the chromatin environment surrounding the promoter that increases promoter availability [17, 37], including nucleosome repositioning in the case of HIV [38]. The rate at which RNAPII and the associated transcriptional machinery binds to the AP state and transitions to the bound promoter (BP) state is the polymerase binding rate (PBR). In the BP state, which is assumed to describe an initiated but paused promoter [39], the rate at which the promoter releases an elongated transcript and returns to the AP state is the polymerase pause release rate (PPRR). Elongated transcripts are translated into protein at a rate Kp, and degradation of mRNA and proteins is modeled as a first-order process. We assume that only one polymerase can bind at a time, and that the polymerase remains paused until released [39, 40]. If the promoter continuously cycles between the AP and BP states, then a burst of transcription occurs, and therefore we refer to this model as a transcriptional cycling model. Alternatively, the BP state returns to the UP state with the same BTR as the AP transitions to UP state, denoting a burst termination event.

**Fig 1.**
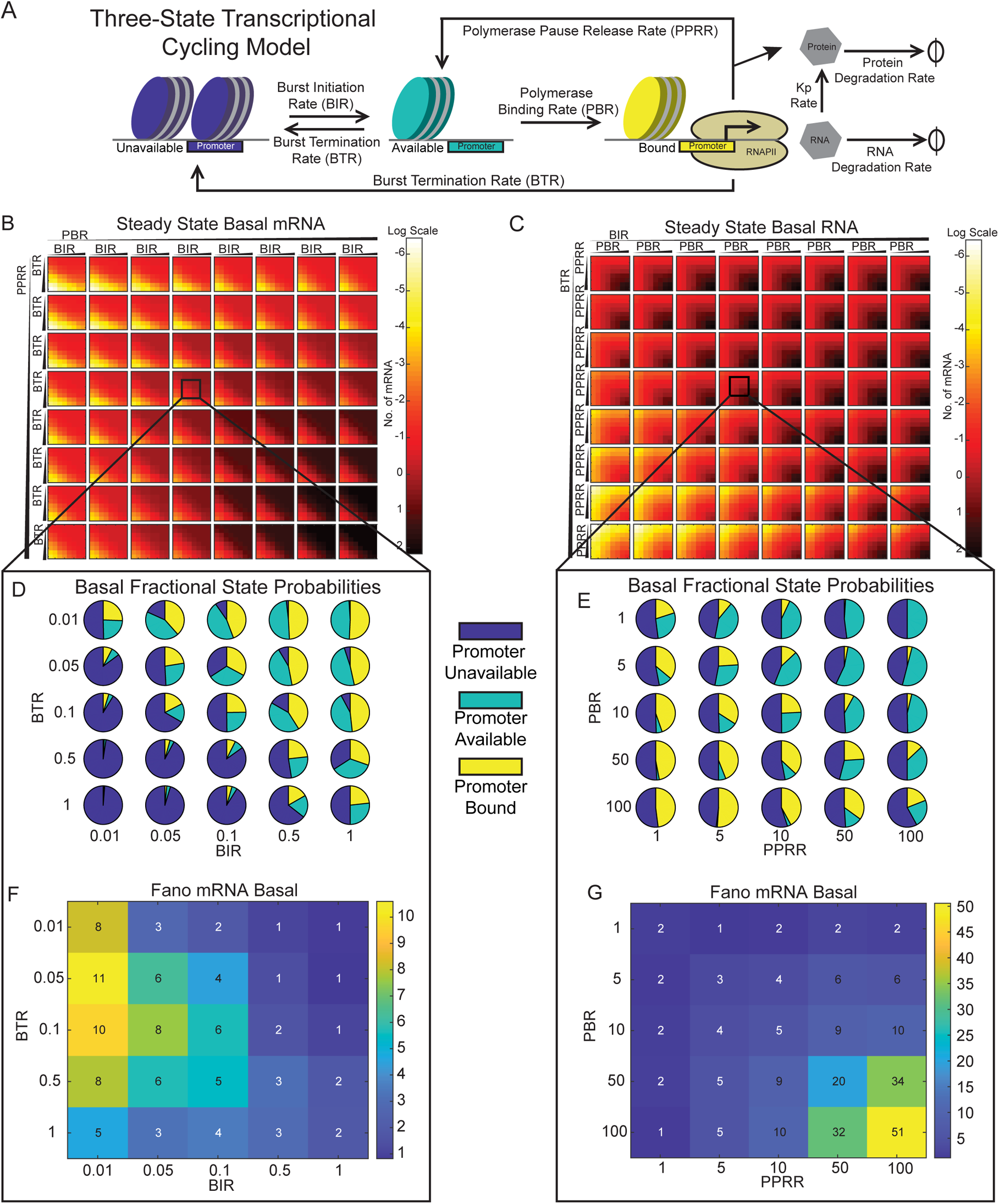
The BIR:BTR ratio controls promoter availability and the PPRR:PBR ratio controls transcriptional cycling. (A) Schematic of three-state transcriptional cycling model, including five species: a bound promoter (BP, in blue), an available promoter (AP, in teal), a bound promoter (BP, in yellow), RNA, and protein. A cycle of transcription occurs when the promoter transitions from BP to AP and back to BP. (B-C) Deterministic solution of steady-state mRNA counts when varying BIR and BTR for fixed values of PBR and PPRR (B) or when varying PBR and PPRR for fixed values of BIR and BTR (C). Parameter ranges are varied low to high via arrow directionality and correspond to the following sets: PBR = [0.1 0.5 1 5 10 50 100 500] hr^−1^, PPRR = [0.1 0.5 1 5 10 50 100] hr^−1^, BIR = [0.005 0.01 0.05 0.1 0.5 1 5 10 50] hr^−1^, and BTR = [0.005 0.01 0.05 0.1 0.5 1 5 10 50] hr^−1^. Heatmap indicates log of average mRNA levels. (D-E) Pie charts representing the fractional probability of UP (blue), AP (teal), and BP (yellow) when varying the BIR:BTR ratio for fixed PBR and PPRR (D) or when varying PBR:PPRR ratio for fixed BIR and BTR (E). (F-G) Fano Factor calculated for the same range of simulations presented in (D) and (E), respectively. Fractional promoter state probabilities and Fano factors were calculated from 1,000 single-cell stochastic simulations under basal conditions out to 10 days for each parameter combination. Square inset in (B) corresponds to the following parameter set: PBR=10 hr^−1^, PPRR=10 hr^−1^, BIR= [0.01 0.05 0.1 0.5 1] hr^−1^, and BTR= [0.01 0.05 0.1 0.5 1] hr^−1^. Square inset in (C) corresponds to the following parameter set: BTR=0.1 hr^−1^, BIR=0.1 hr^−1^, PPRR= [1 5 10 50 100] hr^−1^, and PBR= [1 5 10 50 100] hr^−1^.

This transcriptional cycling model was shown to accurately reflect the biological regulatory mechanisms of burst initiation and pause release for steady-state gene expression [19]. At higher PPRR or PBR values, the exact solution of this model reduces to the two-state model, as the transition from unavailable promoter to available promoter would produce transcripts constitutively [7]. However, it is unclear how variability in inducible transcription would be affected when initiated under different steady-state conditions, which could reflect different chromatin environments.

To explore this, we used HIV as a model system. As a retrovirus, HIV integrates into the infected host cell’s genome [41]. Although most viral integrations lead to productive infections, a rare subset of integrated viruses transition to a latent state, in which little or no virus is actively produced [42]. There is no cure for HIV infection due to this latent reservoir, but quiescent viruses can be reactivated upon stimulation with certain extracellular cues, offering a clinically promising strategy to purge the latent viral reservoir via exploiting the molecular mechanisms that cause activation. The HIV long terminal repeat (LTR) promoter is regulated by NF-κB [43], and TNF stimulation leads to the accumulation of the NF-κB RelA:p50 heterodimer in the nucleus, leading to transcriptional activation [44]. Notably, HIV exhibits bursty transcription that is dependent on the chromatin environment at the location of viral integration [5, 45], with more open chromatin environments leading to higher levels of activation [46, 47].

We first sought to understand the parameter space of the three-state transcriptional cycling model (Fig 1A) by examining average steady-state solutions for HIV mRNA and promoter-state distributions calculated from the five ordinary differential equations (ODEs) in the model (see Materials and Methods). Degradation and translation rates were held constant at experimentally determined values for HIV [18,48,49], while the remaining four parameters were varied over a wide biological range. Examination of the 4-D parameter space revealed regions of varying fractional promoter state probabilities (S1 FigA-C). Higher BTR values increased the probability of promoters residing in the UP state (S1 FigA), whereas increases in BIR resulted in a higher probability of having promoters in the AP and BP states (S1 FigB and C). Increases in PPRR were associated with a higher probability of being in the AP state and increases in PBR increased the probability of being in the BP state given a certain UP probability (BIR to BTR). The clear patterns along the diagonal within each BIR-BTR square strongly suggested that the ratio of these parameters regulates the distribution of promoter states.

To support these qualitative observations, we derived quantitative relationships between parameter ratios and the ratios of promoter states by examining the five ODEs at steady-state. Overall, we found that

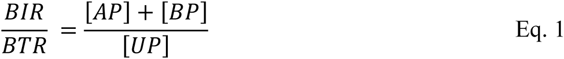

indicating that the relative magnitude of BIR to BTR controls the ratio of promoters in the AP+BP to UP state (Equation **Error! Reference source not found.**). In addition,

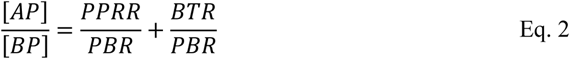

indicating that the relative magnitude of PBR to BTR+PPRR controls the ratio of promoters in the AP to BP state (Equation 2). As seen in Equation 2, the relative strengths of both PPRR to PBR and BTR to PBR toggle the amount of AP state compared to the BP state. Within our chosen parameter space, PPRR is usually greater than BTR (reflecting the different time scales of these biological processes), allowing the ratio of AP states to BP states to simplify to the relative magnitude of PPRR to PBR. For fixed values of BIR and PBR, lower PPRR and higher BTR values lead to low steady-state RNA production (S1 FigD), consistent with the finding that UP dominates the fractional state probabilities (S1 FigE). With information regarding BIR:BTR and PPRR:PBR ratios, we can summarize the initial fractional distributions of the three promoter states, with BIR:BTR controlling promoter accessibility (i.e., the amount of closed promoter relative to open promoter, UP to AP+BP), and PPRR:PBR controlling transcriptional cycling (i.e., the switching between bound and unbound open promoter, AP to BP).

Under steady-state conditions, the fractional distribution of the initial promoter states affects the amount of mRNA produced. We visualized how the BIR:BTR ratio and the PPRR:PBR ratio differentially affect steady-state RNA production using different heatmap configurations. We first examined the effect of the BIR:BTR ratio (Fig 1B) by varying BIR and BTR (inner heatmaps) for a range of fixed values of PBR and PPRR (outer grid). As PBR and PPRR increased, average steady-state mRNA levels increased, with the highest levels at the bottom right corner (Fig 1B). Within each square (i.e., for a fixed value of PBR and PPRR), average mRNA levels varied with the BIR:BTR ratio, as can be observed by the patterning along the diagonal, with average mRNA increasing from low BIR:BTR ratios (bottom left corner) to high BIR:BTR ratios (upper right corner) within each inner heatmap. In our PBR:PPRR heatmaps for fixed values of BIR and BTR (Fig 1C), we observed a different pattern, noting that average mRNA level increased as PBR and PPRR increased (Fig 1C, upper left to bottom right of each heatmap). However, the smaller relative value of PBR versus PPRR acts as a rate-limiting step for the overall transcriptional cycle [7]. Thus, as BIR increases and BTR decreases across the entire grid, average steady-state mRNA levels generally increase, but the variation is limited by the PBR:PPRR ratio (Fig 1C).

To observe the distribution of initial promoter states varied with these ratios, we zoomed in on one inner heat map. First, we fixed PPRR and PBR at values of 10 (i.e., “outer” PPRR:PBR = 1), and then visually explored how the distribution of initial promoter states varied for a range of BIR:BTR ratios (Fig 1B, box). For this range, we observed that the fraction of promoters that exist in the UP state decreased with increasing BIR:BTR (Fig 1D), consistent with Equation 1 and average mRNA level. Note that because the ratio of PPRR:PBR is 1, the non-UP fraction is equally distributed between the AP (teal) and BP (yellow) states. Repeating this analysis for a fixed BIR:BTR ratio of 1 (Fig 1C, box), we observed that varying PPRR:PBR changes the relative amounts of AP and BP, while the total fraction of UP remains approximately constant (at ∼50% for BIR:BTR = 1), consistent with Equation 2 (Fig 1E). We observed the highest fraction of BP in the bottom left corner, and the highest fraction of AP in the top right corner.

Previous studies suggest that variations in chromatin environments generate cellular heterogeneity [17,25,50,51]. Varied chromatin environments can be represented in the transcriptional cycling model by initializing promoters with different BIR:BTR and PPRR:PBR ratios leading to variations in the distribution of initial promoter states, and therefore we were interested in how varying these ratios would affect transcriptional noise. To examine this, we calculated how the Fano factor (*F* = σ^2^/μ) varied for the same “zoomed in” heatmaps. We observed that the BIR:BTR had a modest influence on Fano, with the highest values generally observed when BIR:BTR < 1 (Fig 1F). Notably, a BIR:BTR ratio of less than one generally characterizes bursty transcription processes, for which activation rates are less than inactivation rates [52]. However, for BIR:BTR << 1, Fano decreases because the fractional probability of non-UP becomes negligible, and very few cells produce transcripts (Fig 1F, bottom left corner).

Interestingly, we observed that noise is independent of the PPRR:PBR ratio, as the Fano graph is symmetrical about the diagonal (Fig 1G). Rather, Fano increases as both PPRR and PBR increase because, while the probability of occupying the UP state stays approximately constant, the promoters switching between the AP or BP states produce more mRNA through increased transcriptional cycling, enhancing differences between the closed (UP) and open (AP+BP) states. The rate-limiting nature of the transcriptional cycle in the three-state promoter system requires equal contribution of PBR and PPRR for high levels of transcriptional activity in the basal state. In other words, if only one rate of this cycle increases (for instance, a higher PBR compared to a PPRR), the cycle remains paused. Thus, while PPRR and PBR control amounts of AP and BP respectively, changes in expression and noise are more dependent on the absolute values of PPRR and PBR.

### Activating gene expression under different promoter initialization states affects transcriptional noise and cell-to-cell heterogeneity

A key biological question is how activation of TFs upon exogenous stimulation affects transcriptional noise [27, 28]. TFs often recruit molecular complexes that alter one or more of the rates in the transcriptional cycling model. For example, as discussed above, a consequence of activating NF-κB is to mediate the recruitment of P-TEFb, releasing paused RNAPII and allowing transcriptional elongation to proceed more efficiently [32–34]. Because regulation of RNAPII pause release is a widely conserved mechanism to control transcription [53–56], we simulated this activation pathway by increasing the pause release rate, PPRR, in our model and quantified how it affected transcriptional noise.

We first conducted a sensitivity analysis of how increasing PPRR affected transcription over 24 hours, monitoring mRNA over various combinations of BIR, BTR and PBR. We selected a 2-fold increase in PPRR as an appropriate activation signal (S2 FigA), as this allowed activation from basal conditions (mRNA > 3) without overactivation. Starting from the basal steady-state (i.e., the state after 10-day simulations with basal PPRR), we tracked activation of transcription by running 1,000 stochastic simulations of the model for each parameter condition for 24 hours following a 2-fold increase in PPRR. To capture a wide range of initial fractional promoter states, the four parameters were varied over two orders of magnitude (Table 1).

**Table 1:** Model development and parameter space.

| Reaction | Rate | Value (or<br>Range used) | References |
| --- | --- | --- | --- |
| $UP \rightarrow AP$ | BIR (Burst Initiation Rate) | $[0.01 - 1] \text{ hr}^{-1}$ | [7,52,57–60] |
| $AP \rightarrow BP$ | PBR (Polymerase Binding Rate) | $[1 - 100] \text{ hr}^{-1}$ | [7,52,57,61–63] |
| $AP \rightarrow UP$<br>$BP \rightarrow UP$ | BTR (Burst Termination Rate) | $[0.01 - 1] \text{ hr}^{-1}$ | [7,52,57,58,60] |
| $BP \rightarrow RNA + AP$ | PPRR (Polymerase Pause Release<br>Rate) | $[1 - 100] \text{ hr}^{-1}$ | [7,52,62,63] |
| $RNA \rightarrow Protein$ | Kp (Translation Rate) | 1 | [4] |
| $RNA \rightarrow \emptyset$ | DegR (Degradation of RNA Rate) | 0.34 | [4] |
| $Protein \rightarrow \emptyset$ | DegP (Degradation of Protein Rate) | 0.1 | [4] |

We first examined activation trajectories for promoters initiated with three different BIR:BTR ratios (0.1, 1 and 10) for the same fixed PBR and PPRR as in Fig 1B. Increasing BIR:BTR decreased the fraction of promoters in the UP to the AP+BP states under basal conditions (Fig 2A, left). Upon activation (i.e., a 2-fold increase in PPRR), a fraction of promoter states switched from the BP to AP state, but the fraction of promoters in the UP state remained approximately constant for all three ratios, as increasing PPRR does not affect the UP state (Fig 2A, right). Examining the transcriptional trajectories over 24 hours revealed three distinct behaviors (Fig 2B). Initializing promoters with a low BIR:BTR ratio (BIR:BTR = 0.1) rarely led to transcriptional activation, as evidenced by only a few non-zero trajectories and a probability density function (PDF) containing cells that are mostly off (Fig 2B, top). In contrast, initializing promoters with a high BIR:BTR ratio (BIR:BTR = 10) resulted in almost complete activation (Fig 2B, bottom). Interestingly, initializing with a BIR:BTR ratio of 1 resulted in heterogeneous outcomes, characterized by bimodal activation. As expected, average mRNA increased over 24 hours in all cases, with the largest absolute increase observed for the BIR:BTR ratio of 10 (Fig 2C). Transcriptional noise (as quantified by Fano) also increased over 24 hours in all cases, although both BIR:BTR=10, and BIR:BTR=1 peak at 1 and 4 hours respectively (Fig 2D). This plateau of noisy expression for the higher BIR:BTR ratios reflects the required time necessary for the stimulated system to equilibrate after PPRR increases (Fig 2B, bottom and middle panel). Overall, this illustrates how the different distributions of initial promoter states that result from different initial BIR:BTR ratios lead to varied phenotypes upon activation, with the basal fraction of UP inversely correlated to the fraction of activating cells.

**Fig 2.**
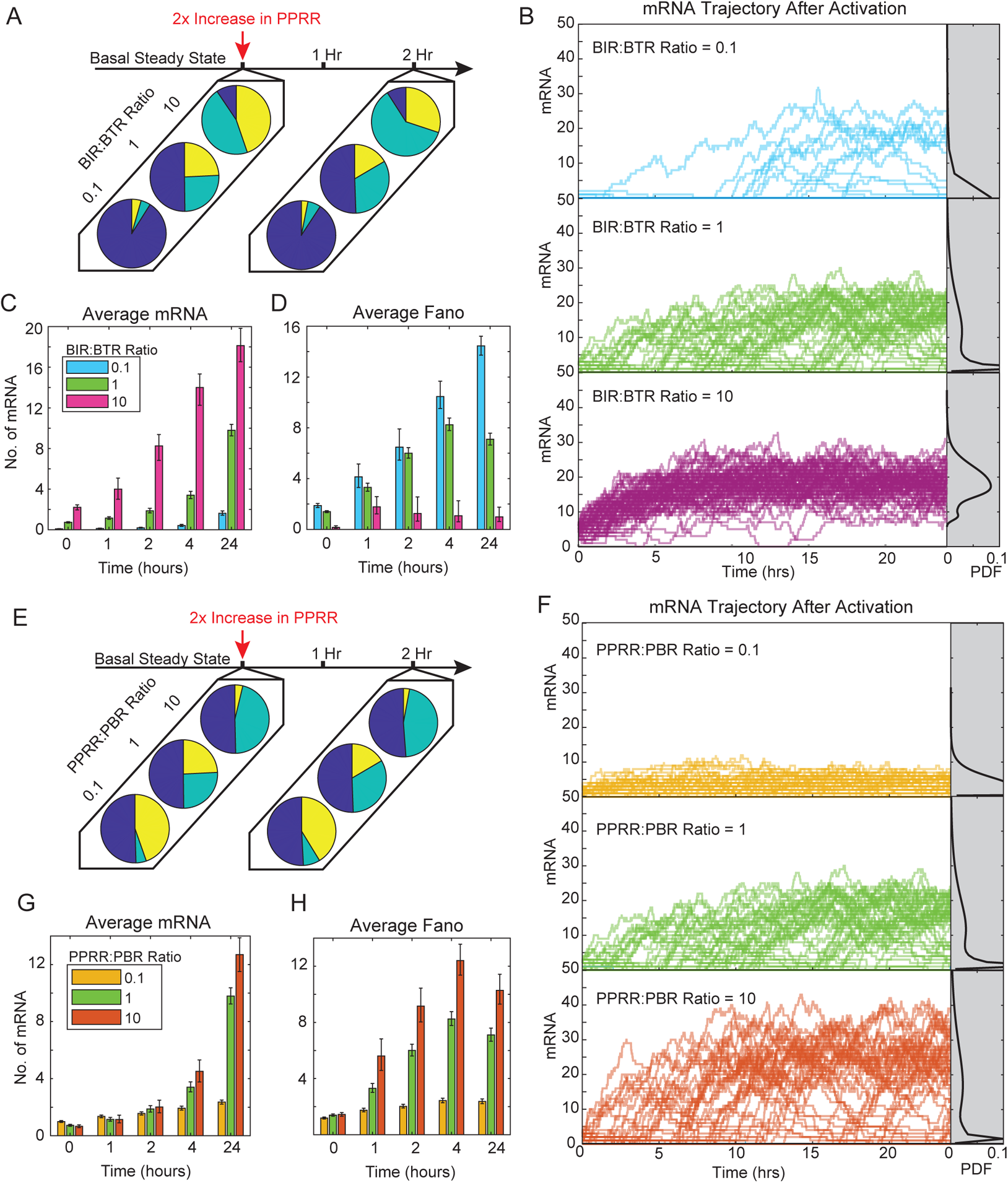
The initial distribution of promoter states influences the heterogeneity in transcriptional activation modeled as an increase in PPRR. (A) Three representative pie charts of fractional promoter-state probability of UP (blue), AP (teal) and BP (yellow) for BIR:BTR ratios of 0.1, 1, and 10 with PBR and PPRR held constant at 10 hr^−1^. Basal conditions were calculated from 1000 single-cell stochastic simulations out to 10 days for each parameter combination. At time=0, PPRR was increased two-fold (B), and new fractional probabilities were captured at 2 hours. (B) Representative trajectories for BIR:BTR = 10 (pink), BIR:BTR = 1 (green), and BIR:BTR = 0.1 (cerulean). Each line represents one stochastic simulation out to 24 hours. Only 100 simulations are plotted for each condition for ease of visualization. Gray regions on the right represent the probability density of mRNA counts at 24 hours, with kernel smoothing. (C-D) Average mRNA counts (C) and Fano factor (D) for the three BIR:BTR ratios at 0, 1, 2, 4 and 24 hours. Average mRNA values and Fano were calculated from 1,000 single-cell stochastic simulation for each parameter combination. Error bars represent 95% bootstrapped confidence intervals. (E) Three representative pie charts of fractional promoter-state probability of UP (blue), AP (teal) and BP (yellow) for PPRR:BBR ratios of 0.1, 1, and 10 with BIR and BTR held constant at 0.1 hr^−1^. Fractional probabilities were calculated as described in (A). (F) Representative trajectories for PPRR:BBR = 0.1 (yellow), PPRR:PBR = 1 (green), and PPRR:PBR = 10 (orange). Data presented as described in (B). (G-H) Average mRNA counts (G) and Fano factor (H) for the three PPRR:PBR ratios at 0, 1, 2, 4 and 24 hours. Data presented as described in (C-D).

Next, we examined how varying the PPRR:PBR ratio (0.1, 1, and 10) contributes to noise and heterogeneity upon PPRR activation. As noted previously, under basal conditions, as the ratio of PPRR:PBR decreases, the fraction of BP increases, which the fraction of UP remains constant (Fig 2E, left). Upon stimulation (i.e., a 2-fold increase in PPRR at time=0), the amount of AP increased marginally by 2 hours (Fig 2E, right). Increasing the PPRR:PBR ratio over 24 hours increased both the fraction of promoters actively transcribing (Fig 2F) and the average mRNA by 24 hours (Fig 2G). This increase in mRNA production is accompanied by increases in noise (Fano; Fig 2H). This increase peaked around the 4-hour mark for PPRR:PBR ratios of 1 and 10, corresponding to when trajectories begin to plateau, and leads to bimodal mRNA levels across cells by 24 hours (Fig 2F, middle and bottom grey). Lower PPRR:PBR ratios lead to lower levels of gene transcription, and decreased Fano levels, as the trajectories cluster near low expression levels (Fig 3F, top). However, while higher PPRR:PBR ratios increased activation, noise also increased, because the UP fraction remained unchanged and thus there were non-activated cells by 24 hours even at the highest PPRR:PBR ratio.

**Fig 3.**
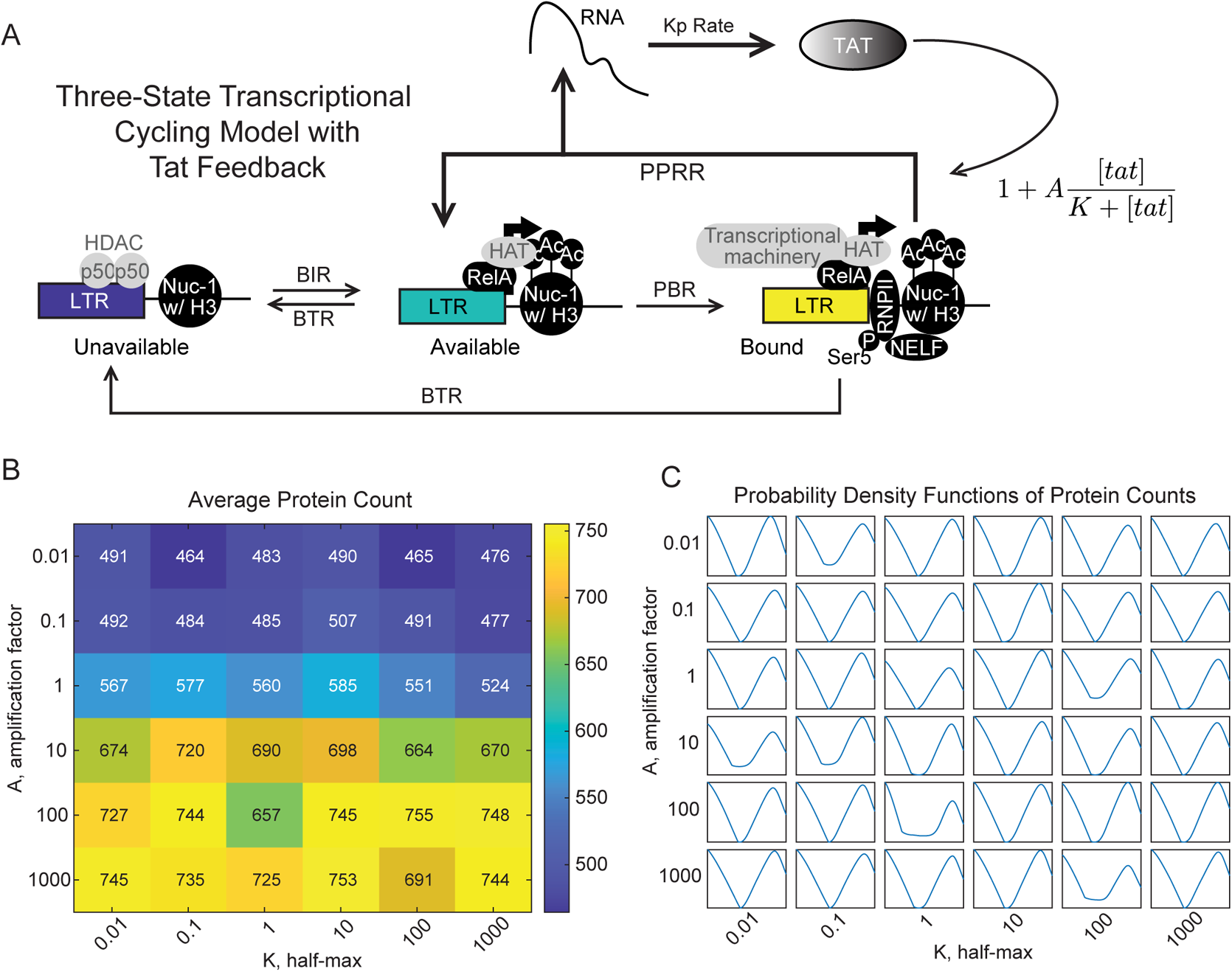
Positive feedback on PPRR activation does not influence bimodality of the mRNA and protein distributions in the three-state transcriptional cycling model. (A) Updated three-state promoter system with HIV nucleosome remodeling, RelA recruitment, and Tat-mediated transcript elongation, which is amplified via positive feedback. Positive feedback is modeled as a saturating function with an amplitude, A, and half-max, K. (B) Heatmap of average protein counts at 24 hours with feedback. Protein counts were generated through stochastic simulation for 1,000 cells for each combination of K and A, which were varied over 5 orders of magnitude. The other parameters were fixed as follows: BIR = 0.1 hr^−1^, BTR = 1 hr^−1^, PBR = PPRR = 10 hr^−1^ (C) Kernel fittings of mRNA counts at 24 hours. Each box contains the probability density curve for that parameter combination.

To generalize our observations of bimodality, we identified a section of the parameter space that we classified as “always on,” “always off,” and “bimodal” (for classification breakdown, see Materials and Methods) (S2 FigB). We then analyzed the initial fractional promoter states for each of these phenotypes (S2 FigC). The “always off” phenotype had much higher levels of UP relative to AP and BP, whereas the “always on” phenotype had approximately equivalent amounts of each promoter. The “bimodal” phenotype was in the middle, with higher UP as compared to the “always on” population, but lower BP and AP as compared the “always off”. This suggests that the distribution of initial promoter states contributes to bimodality following transcriptional activation.

### Positive feedback that amplifies the polymerase pause release rate (PPRR) in the transcriptional cycling model does not affect bimodality

Positive feedback is another common motif used to activate and amplify gene expression [38, 49], and so we next sought to explore how positive feedback would influence gene expression noise following inducible activation. There are many different biological mechanisms of positive feedback, and so we limited our exploration to the mechanism used by HIV. Briefly, upon initiation of HIV transcription, the initiated transcripts form a stem-loop structure referred to as the transactivating response region (TAR), which leads to promoter proximal pausing by RNAPII [64]. HIV encodes its own transcriptional transactivator (Tat), which recruits P-TEFb to the transcriptional start site [65, 66] where it releases paused RNAPII to enable elongation and the generation of a full-length transcript [6,36,67]. HIV exhibits bimodal activation under basal conditions [68] and following stimulation with TNF [69], and therefore we were interested to see to what extent positive feedback contributed to this observation.

We added Tat-mediated positive feedback to the transcriptional cycling model by amplifying PPRR with a Tat-dependent term (Fig 3A) as shown:

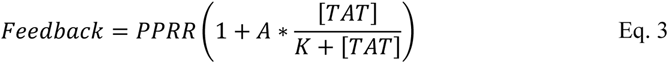

in which K determines the half-maximal saturation and A is the amplitude of the feedback. We performed a sensitivity analysis to determine how these two terms affected transcriptional activation by 24 hours. We chose a region of the parameter space with bursty characteristics (BIR:BTR ratio < 1, and a PPRR:PBR ratio of 1 to remove any rate limiting effects). We found that increasing A by five orders of magnitude only increased average mRNA and protein levels approximately 1.5-fold, while increasing K across a similar range had almost no effect (Fig 3B and S3 FigA). Varying A and K did not affect the fraction of promoters in the UP state (S3 FigB), consistent with the analysis that the UP fraction is unaffected by changes in PPRR. Varying A did affect the fraction of promoters in the AP and BP states, while K had no effect. As the amount of AP saturates the fractional availability, the additional feedback strength will not alter transcriptional production. The influence of K on the sensitivity of the feedback can be visualized by plotting the relative feedback value for varying levels of K. As K decreases, it reaches high feedback strength at very low level of protein production, saturating the signal (S3 FigC). Once saturated, the limiting rate becomes PBR and amplifying PPRR via positive feedback will not lead to further increases in protein production.

Previous studies have suggested that Tat-positive feedback is required for bimodality of protein production [70]. However, we observed that, for our regions of interest in the parameter space, implementing positive feedback as an amplification of PPRR did not alter the bimodality observed in the absence of feedback (Fig 3C). Overall, we conclude that for the transcriptional cycling model in which Tat feedback only affects RNAPII pause release (i.e., PPRR), Tat positive feedback increases overall expression but does not alter bimodality.

### Positive feedback on PPRR increases transcriptional noise when initialized from promoter states with high PPRR:PBR ratios

Although Tat-mediated feedback on PPRR in the transcriptional cycling model does not affect bimodality, it does influence the rate of transcriptional cycling, increasing the rate at which BP transitions back to AP with a release of transcript (Fig 3A). To visualize how feedback affects activation in different regions of the parameter space, we returned to our BIR:BTR and PPRR:PBR ratios to examine mRNA averages, noise profiles, and trajectories after activation. We modeled Tat feedback through the addition of EquationEq. 3, and stochastically simulated our three-state system over an interval of ten days to reach a basal steady-state. The system was then activated by increasing PPRR 2-fold for the same parameter regions explore in Fig 2, but this time in the presence of Tat feedback.

We first examined how feedback affects activation for different BIR:BTR ratios. Although feedback does not affect the initial fraction of promoters in the UP state, it does move more of the non-UP fraction from the BP to the AP state (Fig 4A, left), as the effective PPRR is increased with the addition of feedback. After activation via a 2-fold increase in PPRR, even more of the BP fraction is converted to AP (Fig 4A, right). We saw similar trends in the trajectories of the three BIR:BTR ratios as compared to our results without feedback (Fig 4B; compare to Fig 2B), however they were differentially affected by feedback. For the highest BIR:BTR ratio (BIR:BTR = 10), feedback increased the average mRNA count by approximately 1.4-fold over 24 hours (Fig 4B, top, black versus red, and Fig 4C). By contrast, feedback only moderately increased average mRNA for the intermediate ratio of BIR:BTR = 1, and it had almost no effect on BIR:BTR = 0.1 (Fig 4B, middle and bottom), as most of these promoters remained in the UP state and thus were unaffected by PPRR amplification.

**Fig 4.**
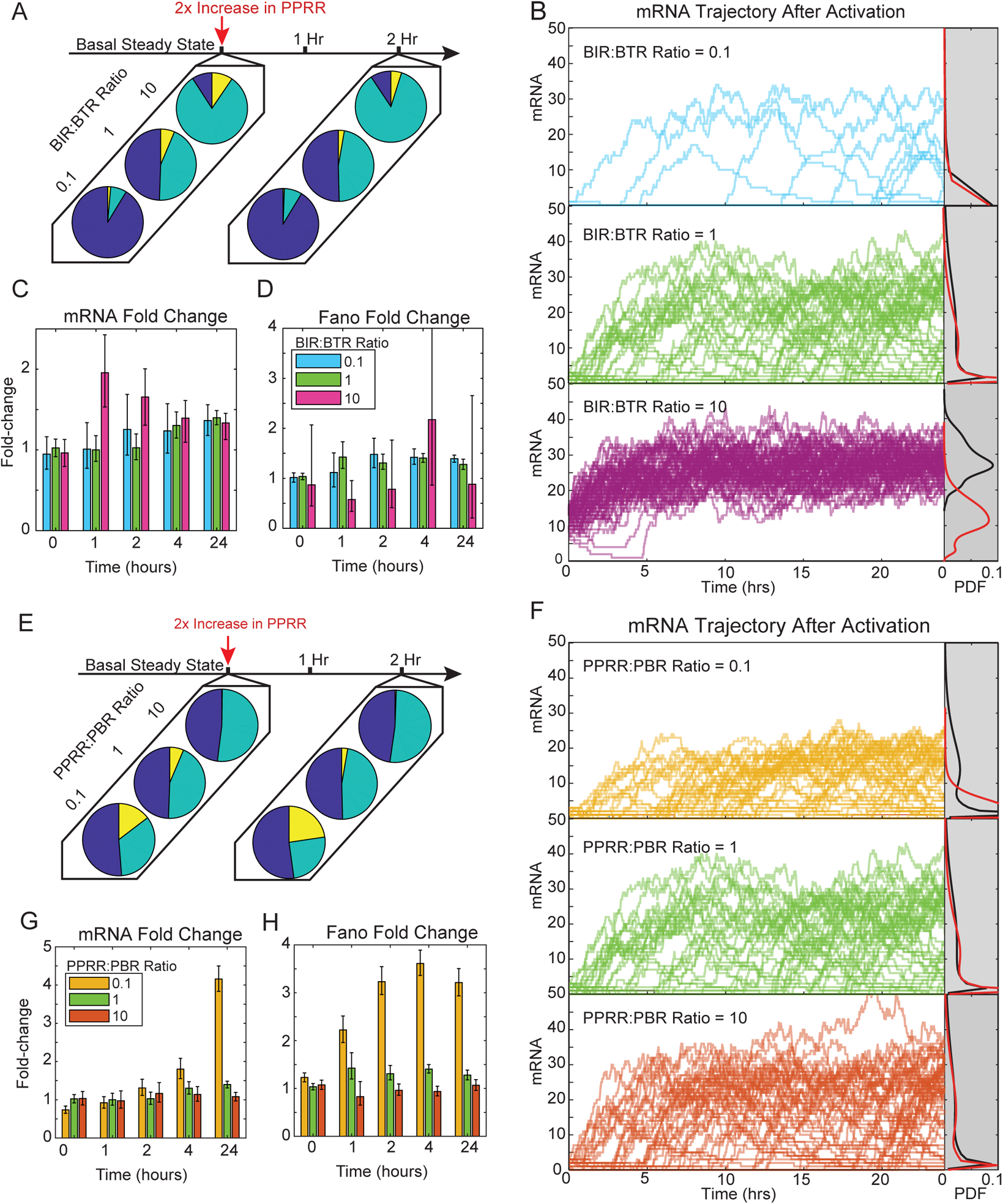
Transcriptional activation in the presence of PPRR positive feedback predominately alters activation of more permissive initial promoter states. (A) Three representative fractional promoter-state probability pie charts with the addition of feedback of UP (blue), AP (teal) and BP (yellow) for BIR:BTR ratios of 0.1, 1, and 10 with PBR and PPRR held constant at 10 hr^−1^. Data presented as described in Fig 2A. (B) Representative simulated trajectories with feedback for BIR:BTR = 10 (pink), BIR:BTR = 1 (green), and BIR:BTR = 0.1 (cerulean) presented as described in (Fig 2B). Gray regions on the right represent the probability density of mRNA counts at 24 hours, with kernel smoothing, with feedback (black) and without feedback (red). (C-D) Fold-change in mRNA counts (C) and Fano factor (D) for the three BIR:BTR ratios at 0, 1, 2, 4 and 24 hours as compared to non-feedback simulations. Data were generated through stochastic simulation for 1,000 cells for each parameter combination. Error bars represent 95% bootstrapped confidence intervals. (E) Three representative pie charts of fractional promoter-state probability with the addition of feedback of UP (blue), AP (teal) and BP (yellow) for PPRR:PBR ratios of 0.1, 1, and 10 with BIR and BTR held constant at 0.1 hr^−1^. Fractional probabilities were calculated as described in (A). (F) Representative simulated trajectories with feedback for PPRR:PBR = 0.1 (yellow), PPRR:PBR = 1 (green), and PPRR:PBR = 10 (orange). Data presented as described in (B). (G-H) Fold-change mRNA counts (G) and Fano factor (H) for the three PPRR:PBR ratios for timepoints of 0, 1, 2, 4 and 24 hours as compared to non-feedback simulations. Data presented as described in (C-D).

Feedback produced only minor increases in noise (i.e., Fano), with the lowest ratio of BIR:BTR=0.1 exhibiting the largest increases by 24 hours (Fig 4D). As expected, feedback played a larger role in amplifying gene transcription for cells that had higher BIR:BTR ratios (and thus less UP). Feedback amplified a few highly active trajectories for BIR:BTR=0.1, but did not alter simulations that were initialized in the UP state, leading to an overall increase in Fano.

Feedback also variably affected activation trajectories initialized across different PPRR:PBR ratios (0.1, 1, and 10; Fig 4E). For the highest ratio, PPRR:PBR = 10, increasing PPRR via feedback did not significantly change the distribution of trajectories, average mRNA levels, or noise over time (Fig 4F-H), because PBR was already the rate limiting step for transcription for this ratio. For the two lower ratios (0.01 and 1), feedback increased both mRNA production and noise over 24 hours (Fig 4G-H). Examining the trajectories, we qualitatively observed a few outlier cells that exhibited large increases in mRNA production with feedback, which likely correspond to the small fraction of cells initialized in the BP state, because feedback amplifies the signal of cells that have available BP. For parameter sets that lead to few cells initiated in the BP state, feedback on PPRR does little to change the phenotype of stimulated cells.

### The three-state transcriptional cycling model qualitatively reproduces experimentally observed activation of latent HIV

Finally, we explored how accurately the transcriptional cycling model could reproduce experimental data on latent HIV activation. We and others have previously used a random telegraph model to describe HIV bursting dynamics [5,18,45,49]. However, the random telegraph model with only two promoter states did not fully the differences in chromatin environment that affected transcriptional bursting of latent viral integrations. For example, for LTR integrations that exhibit similar transcriptional bursting dynamics in the basal state (i.e., are fit by the same basal parameters in the two-state model and have average basal mRNA < 3), we can measure significant differences in the local chromatin environment, as evaluated by the ratio of acetylated histone 3 to total histone 3 (AcH3:H3, Fig 5A). These differences in the basal state are associated with significant differences in transcriptional activation upon stimulation with TNF [4] (Fig 5B). This suggests that the two promoter states of the random telegraph model are insufficient to describe the transcriptional dynamics of these latent HIV integrations in the basal state and during TNF-mediated transcriptional activation. Moreover, stimulation of viral activation using the two-state model did not recapitulate bimodality, a known feature of HIV expression [68, 69]. Therefore, we sought to determine if the transcriptional cycling model could better reflect the biology of HIV reactivation.

**Fig 5.**
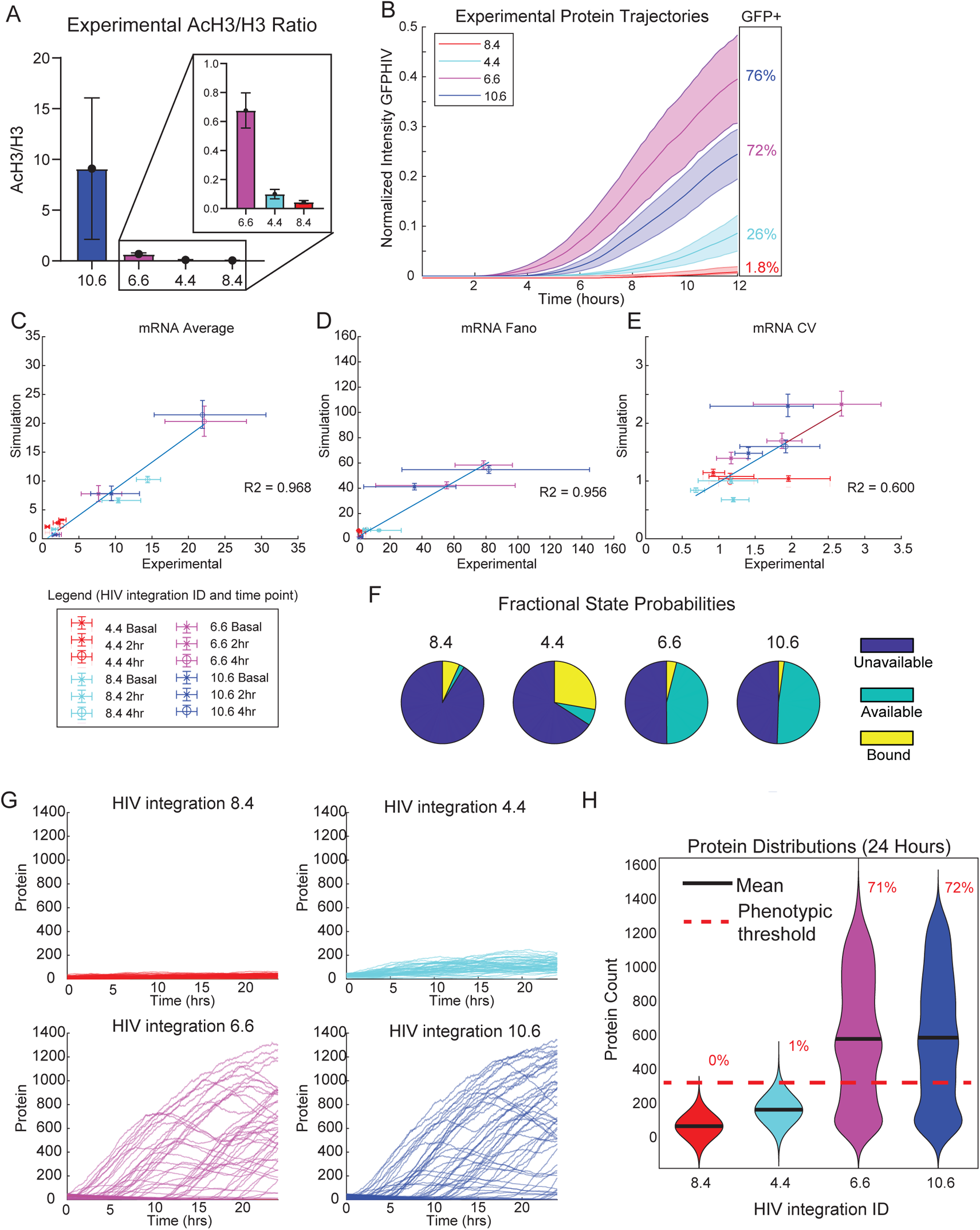
The three-state transcriptional cycling model reproduces transcriptional activation heterogeneity observed for a range of latent HIV integrations. (A) Ratio of enrichment of total histone H3 to acetylated H3 (AcH3) in Jurkat T cells at the indicated target promoters quantified by ChIP-qPCR. Data are presented as mean of % input (non-IP control) ± SD of two biological replicates. (B) Experimental GFP-HIV trajectories for the four clones, plotted with 95% confidence intervals and normalized via experimental setup. (C-E) Scatterplots of three-state promoter simulation with feedback compared to experimental measurements for mRNA Average (C), Fano (D), and CV (E). Error bars represent 95% bootstrapped confidence intervals (C) Fractional state probabilities under basal conditions of UP (blue), AP (teal) and BP (yellow) for the four clones based upon the three-state model. Simulations were run 10,000 times. (F) Representative simulated protein trajectories over 24 hours of the 4 clones. (G) Violin kernel fitting of protein distributions at basal conditions and at 24 hours of the four clones. Black bar represents mean. Red dashed line represents a protein threshold of 275 with percentages as the amount above that threshold. Experimental data in this figure reproduced from [4].

To determine optimal parameter sets describing the four latent HIV integrations, we fit our model to experimental measurements of transcript distributions at 0-, 2- and 4-hours post TNF treatment in four clonal Jurkat T cell populations each with one of the viral integrations [4]. As described above, TNF activation was simulated by increasing PPRR 2-fold. Using a selection algorithm (see Materials and Methods), we found four sets of parameters that minimized error across three features of the experimental measurements of mRNA distributions: the average transcript level, Fano factor, and CV (S4 FigA-C; Table 2).

**Table 2:** Parameters selected for experimental fitting with PPRR activation.

| Parameter | Fitted for<br>Clone 8.4 | Fitted for<br>Clone 4.4 | Fitted for<br>Clone 6.6 | Fitted for<br>Clone 10.6 |
| --- | --- | --- | --- | --- |
| BIR | 1 hr<sup>-1</sup> | 0.5 hr<sup>-1</sup> | 0.1 hr<sup>-1</sup> | 0.1 hr<sup>-1</sup> |
| BTR | 10 hr<sup>-1</sup> | 1 hr<sup>-1</sup> | 0.1 hr<sup>-1</sup> | 0.1 hr<sup>-1</sup> |
| PBR | 50 hr<sup>-1</sup> | 50 hr<sup>-1</sup> | 50 hr<sup>-1</sup> | 50 hr<sup>-1</sup> |
| PPRR | 1 hr<sup>-1</sup> | 1 hr<sup>-1</sup> | 50 hr<sup>-1</sup> | 100 hr<sup>-1</sup> |

We then added Tat positive feedback into the transcriptional cycling model and compared simulation to experimental measurements of transcript distributions in the presence of feedback without further fitting (Fig 5 D-E). We found that average mRNA, Fano, and CV calculated from 1000 simulations reproduced our experimental measurements with feedback with good accuracy (R^2^ values of 0.968, 0.956, and 0.60 respectively), demonstrating that the chosen feedback parameters can reproduce Tat amplification.

When we analyzed the initial promoter states for the selected parameter set for the four integrations, we noted that viral integrations with higher ratios of AcH3:H3 were fit with parameters that exhibited higher ratios of (AP+BP):UP (Fig 5F) but that all were classified as latent (i.e., exhibit average basal mRNA levels < 3). Thus, with the additional parameters to account for chromatin remodeling and polymerase recruitment and release in the three-state model, we identified distributions of initial promoter states that more accurately reflected experimentally measured differences in the chromatin environment.

Finally, we compared our simulations of TNF-activated HIV Tat protein expression over 24 hours to our experimental measurements [4]. We found that increasing PPRR by 2-fold across all four integrations produced significant differences in viral activation by 24 hours. Clones 6.6 and 10.6 exhibited high levels of activation (calculated as 71% and 72% respectively above a basal threshold of 275 simulated proteins, shown via the red dashed line), while 4.4 and 8.4 were not strongly activated (Fig 5G). Moreover, activation of 6.6 and 10.6 was bimodal (Fig 5G), as we had observed experimentally by flow cytometry. While the transcriptional cycling model more accurately captured the chromatin states underlying HIV activation differences, we noticed that the single activation 2-fold increase could not replicate reactivation seen experimentally. Also, the activation of 4.4 did not produce the bimodality seen experimentally. To address these issues, we turned to a multi-step activation model.

### Activation of the transcriptional cycling model via multiple paths reproduces additional observed features of latent HIV activation

As described above, following stimulation by the inflammatory TNF, NF-κB mediates steps in transcriptional activation in addition to its role in the recruitment of P-TEFb [35]. NF-κB also mediates recruitment of CBP/p300 for chromatin remodeling [30], as well as Mediator, RNAPII and other members of the preinitiation complex [31]. Just as we modeled the role of NF-kB in recruiting P-TEFb (i.e., through a 2-fold increase in PPRR), we can recapitulate these additional steps in regulating promoter accessibility and RNAPII binding as fold increases in BIR and PBR, respectively (Fig 6A).

**Fig 6.**
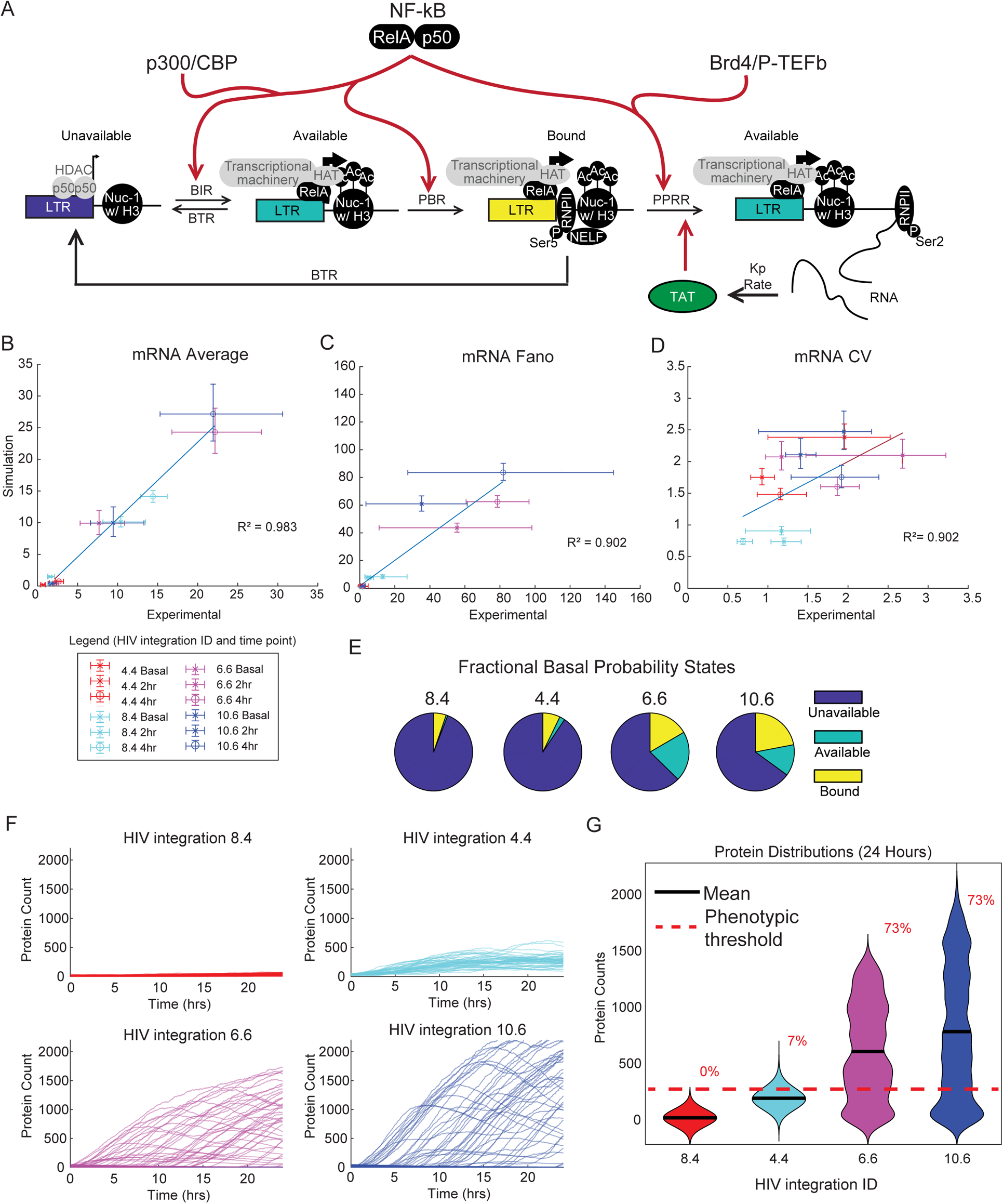
Implementing transcriptional activation via increases in multiple parameters reproduces transcriptional activation heterogeneity observed for a range of latent HIV integrations with more biological accuracy. (A) Schematic of multi-point three-pronged activation in the transcriptional cycling model for HIV. (B-D) Scatterplots of three-state promoter simulation with feedback compared to experimental measurements for mRNA Average (B), Fano (C), and CV (D). Error bars represent 95% bootstrapped confidence intervals. (E) Fractional state probabilities under basal conditions for the four clones based upon the three-state model. Simulations were run 1,000 times. (F) Simulated trajectory protein data of 50 representative cells out to 24 hours of the 4 clones. (G) Violin kernel fitting of protein distributions at basal conditions and at 24 hours of the four clones. Black bar represents the mean. Red dashed line represents a protein threshold of 275 with percentages as the amount above that threshold.

To capture the multi-factorial role of NF-kB activity following TNF stimulation, we increased BIR, PBR, and PPRR 2-fold from basal state conditions across a range of initial parameter values that were consistent with the transcriptional profiles of latent integrations (i.e., average mRNA < 3). Then, using a similar methodology as our PPRR-only activation scheme, we selected a parameter space that most optimally fit the features of transcription at 0, 2 and 4 hours for the experimental measurements of the four latent integrations without feedback (S5 FigA-C). We then ran simulations in the presence of feedback to predict transcript count and distribution at basal, 2, and 4 hours. The parameters chosen as predictions are shown below in Table 3.

**Table 3:** Parameters selected for experimental fitting with multi-point activation

| Parameter | Fitted for<br>Clone 8.4 | Fitted for<br>Clone 4.4 | Fitted for<br>Clone 6.6 | Fitted for<br>Clone 10.6 |
| --- | --- | --- | --- | --- |
| BIR | 0.5 hr<sup>-1</sup> | 0.5 hr<sup>-1</sup> | 0.05 hr<sup>-1</sup> | 0.05 hr<sup>-1</sup> |
| BTR | 10 hr<sup>-1</sup> | 5 hr<sup>-1</sup> | 0.1 hr<sup>-1</sup> | 0.1 hr<sup>-1</sup> |
| PBR | 100 hr<sup>-1</sup> | 100 hr<sup>-1</sup> | 50 hr<sup>-1</sup> | 100 hr<sup>-1</sup> |
| PPRR | 1 hr<sup>-1</sup> | 5 hr<sup>-1</sup> | 10 hr<sup>-1</sup> | 10 hr<sup>-1</sup> |

These predictions for mRNA average, CV, and Fano showed good fit with our experimental dataset in the presence of feedback with an *R*^2^ > 0.90 (Fig 6B-D). The fractional distribution of initial states found to be optimal for these clonal integration positions under basal and latent conditions changed when activation increased all three parameters. All viral integrations had higher levels of UP as compared to the previous simulation (Fig 6E), which is consistent with an activating transcription factor that can act on closed, unavailable chromatin, as NF-kB is known to do. In addition, these parameter sets better capture bursty parameter spaces, where BIR < BTR for all four integrations (Table 2). Jurkat cells harboring viral integrations 6.6 and 10.6 again exhibited bimodal protein distributions by 24 hours (Fig 6F-G), exhibiting increases in activation above threshold to 73%. 4.4 also saw activation to 7% above threshold, suggesting that this multi-activation model better captures reactivation for these more repressed viral integrations. Further optimization might be achieved through reparameterization of feedback terms or fine-tuning of fold-change activations. Altogether, these results demonstrate how implementing multiple TF activation paths in our three-state model more accurately replicates NF-kB-mediated activation of latent HIV viruses.

### The upstream activation path of the transcriptional cycling model differentially affects features of noisy inducible transcription

To compare how each ‘activation path’ contributed to multi-step activation, we took the fitted parameter combinations for the four viral integrations, and examined how BIR, PBR, and PPRR fold-change activation alone affected noise as compared to a multi-step activation. While this parameter space may not replicate our observed experimental data, it allowed for direct comparison across low and high activating viral integrations. For every activation pathway, BIR, PBR, PPRR, or Multi (activating BIR, PBR, and PPRR simultaneously), the appropriate pathway parameter(s) were increased 2-fold and simulated for 24 hours.

Examining transcripts and protein counts at basal and 24 hours post activation (S6 FigA and Fig 7A), we see similar patterns across all viral integrations, with the Multi pathway producing the most transcript and proteins out to 24 hours whatever the initial starting state. However, noise (as quantified by Fano factor) is highest for the low-activating 8.4 and 4.4 viral integrations when activated via the PPRR pathway, while it is highest for 6.6 and 10.6 when activated via the PBR pathway (S6 FigB and Fig 7B). Notably, activation via the BIR pathway leads to a marked decrease in Fano because it increases the frequency of switching between the open and closed promoter states, leading to less separation between the productive and unproductive populations.

**Fig 7.**
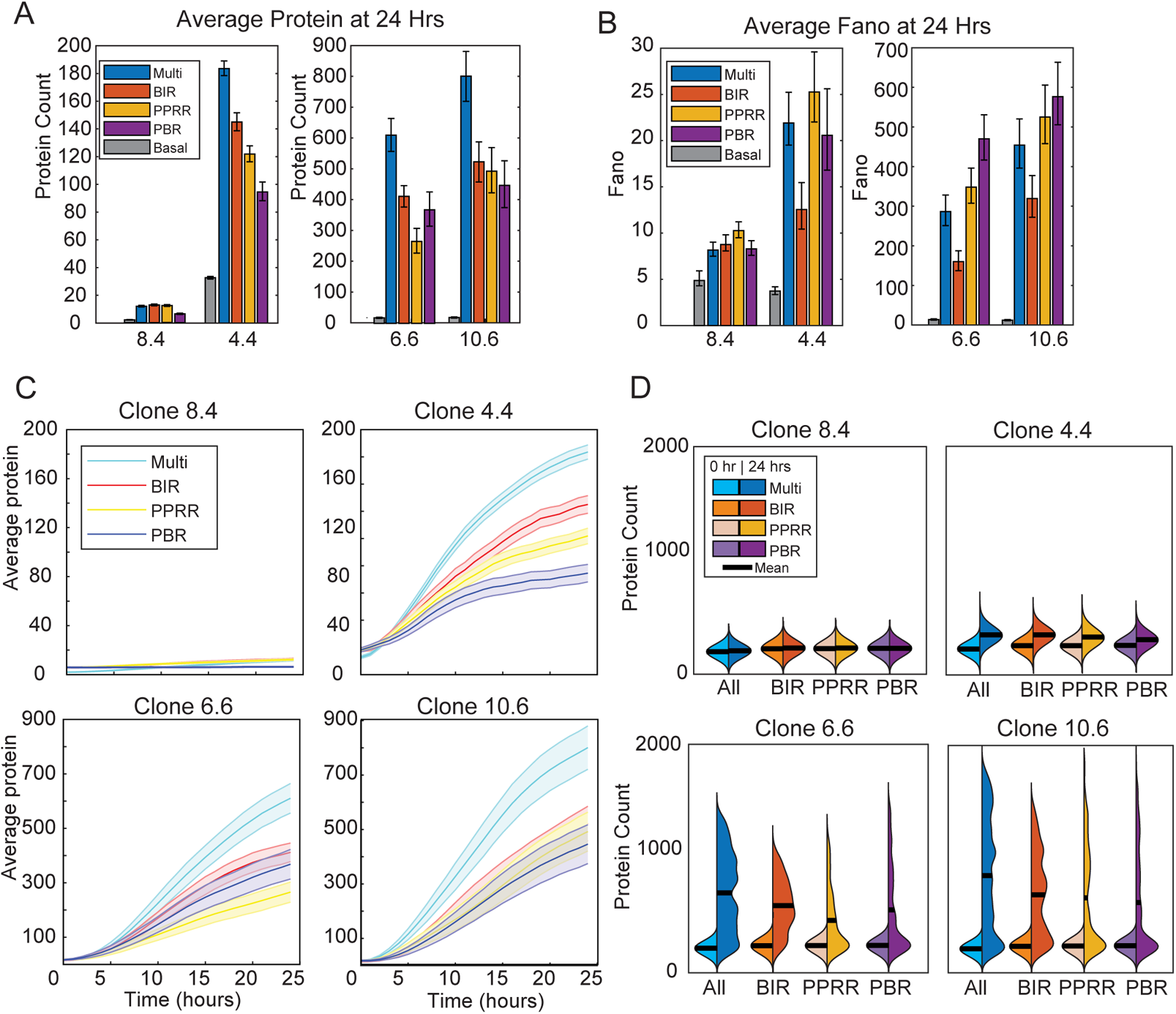
Multi-step activation can be broken down into discrete activation lines which influence noise and protein counts by 24 hours. (A-B) Average protein counts (A) and Fano (B) for the four activation options at 24 hours. Protein counts were generated through stochastic simulation for 1,000 cells for each parameter combination. Error bars represent 95% bootstrapped confidence intervals. (C) Simulated trajectory protein data of 50 representative cells out to 24 hours of the 4 activation lines with the 4 clones. (D) Violin kernel fitting of protein distributions at basal conditions and at 24 hours of the 4 activation lines with the 4 clones. Black bar represents the mean.

To visualize these differences in noise, we plotted the averaged protein trajectories along with kernel probability density functions for each activation pathway (Fig 7C-D). The variations across trajectories for each viral integration highlighted the basal configuration (Fig 6E) upon activation, with 4.4’s changes in BIR leading to increases in simulated protein (Fig 7C top right), as this promotes UP to AP+BP changes. For 6.6 and 10.6, the multi-point activation overall saw the highest increases, with no clear delineation among the individual activation lines. We also compared final protein counts at 24 hours as compared to basal conditions (Fig 7D). Increases to PPRR and PBR for 6.6 and 10.6 lead to more skewed distributions (Fig 7D bottom), with the highest noise but lower activation (S6 FigC). From these comparisons, we can conclude that BIR activation leads to lower noise across clones, without substantial changes to protein average. In contrast, PPRR and PBR increase noise through a few highly productive cells, leading to more skewed distributions. Overall, we conclude that phenotypic heterogeneity following activation by exogenous stimulation depends on the distribution of the initial promoter states, as well as the biological activation pathway.

## Discussion

Despite widespread observations of transcriptional bursting, the molecular mechanisms regulating bursting remain unclear. While experimental studies have suggested a variety of contributors, including RNAPII pausing and chromatin remodeling, the canonical mathematical model of transcriptional bursting is too simple to explore these mechanisms using simulations, and it restricts integrating models of transcriptional bursting with specific processes affected by upstream signals and TFs.

Here, we used a three-state transcriptional cycling model [19] to explore how chromatin variations, polymerase initiation and pausing contribute to transcriptional bursting. We found that the variation in the fractional promoter state probabilities produced by this three-state transcriptional cycling model more accurately described the range of promoter configurations generated by epigenetic variations at basal state. The motivation for this work was to represent the varied chromatin environments at NF-κB-inducible promoters, which correspond to altered transcriptional expression and noise, as we highlighted in recent work [4, 28]. Indeed, we found that these varied promoter state fractions could be used to represent chromatin environments of quiescent-but-inducible promoters in a range of biological contexts (Fig. 1), and to capture the observed differences in noise upon upstream signal activation (Fig. 2).

Our previous studies with endogenous genes demonstrated that target genes with distinct chromatin environments respond differently to TNF stimulation, resulting in different bursting dynamics and ultimately different patterns of inducible gene expression noise [28]. We showed that by increasing histone acetylation via treatment with the HDAC inhibitor trichostatin A (TSA), we could lower gene activation noise. This result is consistent with a more recent and experimentally elegant study demonstrating that increasing histone acetylation via targeted recruitment of the histone acetyltransferase p300 similarly decreased noise in gene expression following TF stimulation [71]. Our model experimentally reproduces this result (Fig. 7), while also allowing for exploration of other effects, such as p300 interactions with Brd4, affecting pause release [72]. Moreover, we have previously reported that transcriptional output is strongly correlated to the fold-change in upstream NF-kB signal activation in individual cells, regardless of initial chromatin state [73]. The role of NF-kB in modulating both BIR:BTR to increase activating fraction, and PPRR:PBR to increase cycling provides a means to explain these findings. Thus, this transcriptional cycling model will be very useful for predicting general dynamics of transcriptional activation and noise for those genes in which chromatin remodeling and polymerase pause release are required steps.

When we applied the three-state model coupled to positive feedback to simulate activation of latent-but-inducible HIV viral integrations, we found that it better represented our experimental observations of chromatin remodeling and RNAPII pause release [4]. Unlike the two-state random telegraph model, the transcriptional cycling model reproduced experimentally observed bimodal patterns of HIV expression within our parameter range. Furthermore, the range of initial promoter states captured by the transcriptional cycling model led to different patterns of gene expression noise following activation (i.e., a sudden change in a model parameter that increases the rate of transcription). This model result was consistent with our experimental observation that the same activating TF can influence multiple mechanisms of activation for our viral integrations, resulting in varied activation profiles. The results of this study highlight the importance of including chromatin remodeling within transcriptional bursting models, particularly when comparing target genes with large variations in initial promoter state fractions, representing different chromatin environments.

Several studies have demonstrated that Tat positive feedback is a direct contributor to bimodal HIV gene expression [68, 70]. Our model, on the other hand, found parameter regions of bimodality that did not require Tat positive feedback, suggesting that the initial promoter state can also encode bimodal responses. In this case, Tat positive feedback amplifies these differences, further separating activated cells from non-activated cells. In settings with more homogeneous chromatin environments, feedback may be required to for bimodality, but we did not explore these parameter regions in this study.

In our study, we used error minimization to choose initial parameter sets for our experimentally measured viral integrations, however these parameter sets are not unique. A limitation of our study is that there are many different parameter sets that would produce qualitatively similar results. Additional work would be required to fit the transcriptional cycling model to experimental data to identify unique parameter sets describing each viral integration. In particular, fitting the non-Poisson moments for RNA and protein distributions might provide additional information to further restrict the possible solutions for initializing the three-state transcriptional cycling model [7, 74]. This could then be used in conjunction with techniques for fitting datasets, such as Bayesian parameter estimation, which requires the knowledge of appropriate priors from the solutions.

Future work with this model allows for studies of how exogeneous perturbations affecting chromatin remodeling combined with other transcriptional regulatory mechanisms (e.g. RNAPII pause release or transcription factor recruitment) synergize to activate gene expression. For example, using TSA to induce chromatin remodeling at the LTR promoter is a clinical strategy to reactivate latent HIV reservoirs to clear the infection [4], and this has been found to synergize with TNF and other NF-κB activators [4,75,76]. Using this model, we can more accurately simulate the effects of multiple drug treatments affecting these pathways and predict how they will impact gene expression noise and viral activation and provide possible avenues for future developments.

## Materials and Methods

### Model Development

The three-state transcriptional cycling model without activation can be represented by the following mass-action kinetic equations:

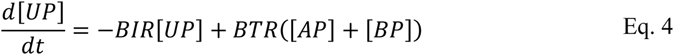

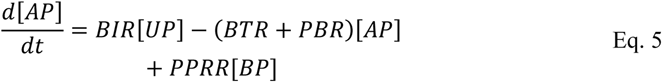

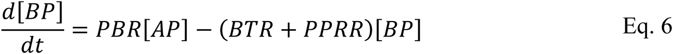

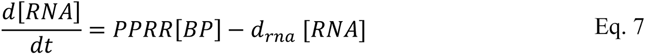

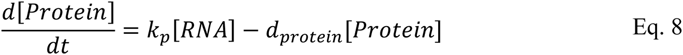

While transcription is stochastic at the single-cell level, we will assume that gene expression follows a deterministic trajectory under population dynamics. We will also assume that the promoter can exist in one of three states, UP, AP, or BP. This conservation of states can be expressed by the following equation.

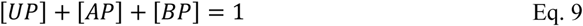

These equations can be deterministically solved using an ODE solver. We utilized the ODE solver in NFSIM [77] to model our system at steady-states conditions a time interval of ten days, capturing dynamics at equal time increments. We used a parameter set that could capture the wide range of mRNA responses, PBR=[0.1 0.5 1 5 10 50 100 500] hr^−1^, PPRR=[0.1 0.5 1 5 10 50 100] hr^−1^, BIR=[0.005 0.01 0.05 0.1 0.5 1 5 10 50] hr^−1^, and BTR=[0.005 0.01 0.05 0.1 0.5 1 5 10 50] hr^−1^. Data was then analyzed via MATLAB.

### Stochastic Simulation and Activation

To stochastically simulate the model, we utilized the SSA utility of NFSIM [77], which implements Gillespie algorithm [78]. The model was first initialized under steady-state basal conditions for ten days. The system was then activated by increasing PPRR two-fold at time = 0, and then simulated out for 24 hours. The promoter states, along with RNA and protein counts were captured in equal time increments. Unless otherwise noted, each parameter combination was stochastically simulated 1,000 times. To classify these simulations, we first tested deviance from unimodality of the probability density distribution utilizing the Hartigan’s dip test [79, 80], with dip > 0.05, and calculating the p value null hypothesis of unimodal distribution > 0.15.

Simulations that met these criteria were classified as “bimodal.” For the remainder of the simulations, we separated “always on” from “always off” with a threshold value of 250 proteins, comparable to thresholds shown in a similar to delineate between activated and non-activated cells [49].

We analyzed the distributions for bimodality using At each relevant time point, mRNA and protein averages, and mRNA Fano were calculated, with 95% confidence intervals calculated via bootstrapping (n=10,000 and α = 0.05) as described previously [28]. Trajectories, heatmaps, bar plots, pie charts, violin plots, and kernel estimations were then generated in MATLAB. Other simulations involving protein feedback and multi-activation steps utilized the same parameter combinations and analysis.

To visualize the differences along ratios, six different parameter combinations were selected. Three represented variations in BIR:BTR (as 0.1, 1, and 10), while the other three represented variations in PPRR:PBR (0.1, 1, and 10). Parameter combinations are shown below in Table 4.

**Table 4.**

| Name | BIR | BTR | PBR | PPRR |
| --- | --- | --- | --- | --- |
| BIR:BTR = 0.1 | 0.01 hr<sup>-1</sup> | 0.1 hr<sup>-1</sup> | 10 hr<sup>-1</sup> | 10 hr<sup>-1</sup> |
| BIR:BTR = 1 | 0.1 hr<sup>-1</sup> | 0.1 hr<sup>-1</sup> | 10 hr<sup>-1</sup> | 10 hr<sup>-1</sup> |
| BIR:BTR = 10 | 1 hr<sup>-1</sup> | 0.1 hr<sup>-1</sup> | 10 hr<sup>-1</sup> | 10 hr<sup>-1</sup> |
| PPRR:PBR = 0.1 | 0.1 hr<sup>-1</sup> | 0.1 hr<sup>-1</sup> | 10 hr<sup>-1</sup> | 1 hr<sup>-1</sup> |
| PPRR:PBR = 1 | 0.1 hr<sup>-1</sup> | 0.1 hr<sup>-1</sup> | 10 hr<sup>-1</sup> | 10 hr<sup>-1</sup> |
| PPRR:PBR = 10 | 0.1 hr<sup>-1</sup> | 0.1 hr<sup>-1</sup> | 10 hr<sup>-1</sup> | 100 hr<sup>-1</sup> |

### Clone parameter fitting

We hypothesized that this three-state model can better reflect the biological re-activation of latent HIV. To confirm this hypothesis, we sought to find the particular parameter space that describes the clonal differences present. This parameter space is defined by several assumptions and biological considerations.

The first consideration is for bursting transcription, defined as the infrequent transition from low levels of RNA production to high levels of RNA production. In our model, this reflects the transition from a closed chromatin environment to a more open chromatin environment, and can be represented as the parameter region where,

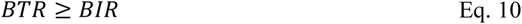

Given our understanding of steady-state analysis, this would result in higher levels of BP as compared to AP and BP.

The second consideration is the differences between our low and high activating clones. The high activating clones see higher levels of acetylation under basal conditions, but the low activating clones see higher levels of histone wrapping. To correlate this to our three-state promoter model, the low activating clones should see higher levels of UP as compared to our high activating clones. We know that the amount of UP us controlled by our BIR to BTR ratio, so we can represent the parameter space through the following inequality:

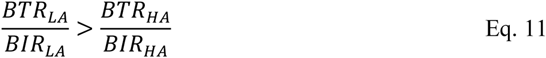

Our third consideration required examining the number of initialized states for the high activating clones. These clones saw higher levels of open chromatin, without significant changes to the amount of transcriptional machinery priming. We can summarize this in our model as AP is greater for high activating clones, whereas the amount of BP remains the same. Given our understanding of steady-state values relating PBR to PPPR, we can reflect this biological consideration through the following inequality:

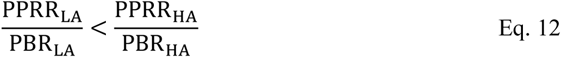

With these considerations, we then sought to find the parameter space that best matched experimental results. Stochastic simulations were run over a biologically plausible parameter space, where basal mRNA > 3 as determined through deterministic solutions and bimodality was observed according to our bimodal criteria. This space was defined as given in Table 1. For each parameter combination, 1000 stochastic simulations were run, with promoter states, mRNA, and protein data captured out to 24 hours. We then used the three considerations above to select appropriate parameter spaces for each clone. To quantify the match, we performed root mean square error (MSE) minimization over mRNA transcript counts, as well as Fano and CV for basal, 2, and 4 hours for the four clones. The best fits for each clone, defined with the lower MSE, were then analyzed, with mRNA counts and noise measurements calculated. Trajectories were graphed with bootstrapped 95% confidence intervals.

### Code and Software Accessibility

NFSIM (v1.11) [77] was used for deterministic and stochastic simulations, with the model written in BNGL (v2.8) [81]. Data was analyzed and plotted in MATLAB (2019b). All code can be found at: https://github.com/miller-jensen-lab/bullock_3state_2022

## Supplemental Figure Legends

**S1 Fig. Steady-state heatmaps and PPRR:BTR comparisons (related to Fig 1).**

(A-C) Heat maps of the deterministic steady state solutions for UP, AP, and BP fractional probabilities. Parameter ranges are represented low to high via arrow directionality and correspond to the following sets: PBR=[0.1 0.5 1 5 10 50 100 500] hr^−1^, PPRR=[0.1 0.5 1 5 10 50 100] hr^−1^, BIR=[0.005 0.01 0.05 0.1 0.5 1 5 10 50] hr^−1^, and BTR=[0.005 0.01 0.05 0.1 0.5 1 5 10 50] hr^−1^. (D) Deterministic solution of MRNA counts representing steady state values plotted as a log scale heatmap. Parameter ranges are represented low to high via arrow directionality and correspond to the following sets: PBR=[0.1 0.5 1 5 10 50 100 500] hr^−1^, PPRR=[0.1 0.5 1 5 10 50 100] hr^−1^, BIR=[0.005 0.01 0.05 0.1 0.5 1 5 10 50] hr^−1^, and BTR=[0.005 0.01 0.05 0.1 0.5 1 5 10 50] hr^−1^. Square inset corresponds to the following parameter set: PBR = 10 hr^−1^, BIR = 0.1 hr^−1^, PPRR = [0.5 1 5 10 50] hr^−1^, and BTR=[0.01 0.05 0.1 0.5 1] hr^−1^. (E-F) Pie charts of the fractional promoter-state probability (E) and Fano factor (F) at each parameter combination. Values determined by stochastic simulation under basal conditions out to 10 days for 1,000 cells for each parameter combination. Promoter states denoted as UP (blue), AP (teal), and BP (yellow).

**S2 Fig. Classification of phenotypes after activation via PPRR increase (related to Fig 2).**

(A) Fold Change mRNA trajectories for PPRR increases of 0.5 (green), 2 (cerulean), 5 (red), 10 (blue), and 50 (purple). Shaded areas are 95% confidence intervals. PPRR and PBR were set at 10 hr^−1^. (B) Average Protein counts at 24 hours. Protein counts were generated through stochastic simulation for 1,000 cells for each parameter combination. Phenotypes of “Always On,” “Bimodal,” and “Always Off” were established by separating out the bimodal population (defined by Hartigan’s Dip Value > 0.15 and a p-test < 0.05 (See Materials and Methods for more information). (C) Bar chart representing the percentage of each promoter state from basal initialization conditions, separated by ending phenotype classification. Error bars represent 95% bootstrapped confidence intervals.

**S3 Fig. Feedback strength influence on fractional state probabilities and protein counts (related to Fig 3).**

(A-B) Heatmap of average mRNA at 24 hours (A) and fractional promoter-state probabilities in the initial state (B) for a range of feedback strengths. All other parameters are set to BIR = 0.1 hr^−1^, BTR = 1 hr^−1^, PBR = PPRR = 10 hr^−1^. Data was generated by stochastic simulation for 1,000 cells for each parameter combination. The feedback terms K (half max) and A (amplification factor) were varied over 5 orders of magnitude. (C) Feedback strength calculated for varied K values and plotted versus protein.

**S4 Fig. Model fits of experimental HIV data compared to simulations using computational PPRR activation scheme (related to Fig 5).**

(A-C) Scatterplots of results from simulation of the three-state transcriptional cycling model without feedback compared to experimental measurements of mRNA distributions from populations of T cells harboring latent HIV integrations and stimulated with TNF with Tat feedback blocked (from ref. 4). Correlation between simulation and experimental data is shown for mRNA average (A), Fano (B), and CV (C). Error bars represent 95% bootstrapped confidence intervals.

**S5 Fig. Model fits of experimental HIV data compare to simulations using computational multi-step activation scheme (related to Fig 6).**

(A-C) Scatterplots of results from simulation of the three-state transcriptional cycling model without feedback compared to experimental measurements of mRNA distributions from populations of T cells harboring latent HIV integrations and stimulated with TNF with Tat feedback blocked (from ref. 4). Correlation between simulation and experimental data is shown for mRNA average (A), Fano (B), and CV (C). Error bars represent 95% bootstrapped confidence intervals.

**S6 Fig. Comparisons of average mRNA and Fano factor across activation paths (related to Fig 7).**

(A-B) Average mRNA counts (A) and Fano factor (B) for the four activation paths across the four HIV integrations for timepoints of 0, 1, 2, 4 and 24 hours. mRNA counts were generated through stochastic simulation for 1,000 cells for each parameter combination. Error bars represent 95% bootstrapped confidence intervals. (C) Percentage of cells above ‘ON’ threshold (set at 250 proteins per cell) at 24 hours.

## References

1. Limi S, Senecal A, Coleman R, Lopez-Jones M, Guo P, Polumbo C, et al. Transcriptional burst fraction and size dynamics during lens fiber cell differentiation and detailed insights into the denucleation process. J Biol Chem. 2018;293: 13176–13190. doi:10.1074/jbc.RA118.001927

2. Tunnacliffe E, Corrigan AM, Chubb JR. Promoter-mediated diversification of transcriptional bursting dynamics following gene duplication. Proc Natl Acad Sci. 2018;115: 8364–8369. doi:10.1073/pnas.1800943115

3. Dar RD, Shaffer SM, Singh A, Razooky BS, Simpson ML, Raj A, et al. Transcriptional Bursting Explains the Noise–Versus–Mean Relationship in mRNA and Protein Levels. Chauhan A, editor. PLoS One. 2016;11: e0158298. doi:10.1371/journal.pone.0158298

4. Wong VC, Bass VL, Bullock ME, Chavali AK, Lee REC, Mothes W, et al. NF-κB-Chromatin Interactions Drive Diverse Phenotypes by Modulating Transcriptional Noise. Cell Rep. 2018;22: 585–599. doi:10.1016/j.celrep.2017.12.080

5. Singh A, Razooky BS, Cox CD, Simpson ML, Weinberger LS. Transcriptional Bursting from the HIV-1 Promoter Is a Significant Source of Stochastic Noise in HIV-1 Gene Expression. Biophys J. 2010;98: L32–L34. doi:10.1016/j.bpj.2010.03.001

6. Tantale K, Garcia-Oliver E, Robert M-C, L’Hostis A, Yang Y, Tsanov N, et al. Stochastic pausing at latent HIV-1 promoters generates transcriptional bursting. Nat Commun. 2021;12: 4503. doi:10.1038/s41467-021-24462-5

7. Cao Z, Filatova T, Oyarzún DA, Grima R. A Stochastic Model of Gene Expression with Polymerase Recruitment and Pause Release. Biophys J. 2020;119: 1002–1014. doi:10.1016/j.bpj.2020.07.020

8. Shaffer SM, Dunagin MC, Torborg SR, Torre EA, Emert B, Krepler C, et al. Rare cell variability and drug-induced reprogramming as a mode of cancer drug resistance. Nature. 2017;546: 431–435. doi:10.1038/nature22794

9. Sanchez A, Golding I. Genetic Determinants and Cellular Constraints in Noisy Gene Expression. Science (80-). 2013;342: 1188–1193. doi:0.1126/science.1242975

10. Dar RD, Razooky BS, Singh A, Trimeloni T V., McCollum JM, Cox CD, et al. Transcriptional burst frequency and burst size are equally modulated across the human genome. Proc Natl Acad Sci U S A. 2012;109: 17454–9. doi:10.1073/pnas.1213530109

11. Suter DM, Molina N, Gatfield D, Schneider K, Schibler U, Naef F. Mammalian Genes Are Transcribed with Widely Different Bursting Kinetics. Science (80-). 2011;332: 472–474. doi:10.1126/science.1198817

12. Wang DC, Wang X. Clinical significance of spatiotemporal transcriptional bursting and control. Clin Transl Med. 2021;11: e518. doi:10.1002/ctm2.518

13. Senecal A, Munsky B, Proux F, Ly N, Braye FE, Zimmer C, et al. Transcription Factors Modulate c-Fos Transcriptional Bursts. Cell Rep. 2014;8: 75–83. doi:10.1016/j.celrep.2014.05.053

14. Li C, Cesbron F, Oehler M, Brunner M, Höfer T. Frequency Modulation of Transcriptional Bursting Enables Sensitive and Rapid Gene Regulation. Cell Syst. 2018;6: 409–423.e11. doi:10.1016/j.cels.2018.01.012

15. Cavallaro M, Walsh MD, Jones M, Teahan J, Tiberi S, Finkenstädt B, et al. 3 ′-5 ′ crosstalk contributes to transcriptional bursting. Genome Biol. 2021;22: 56. doi:10.1186/s13059-020-02227-5

16. Zhao C, Mirando AC, Sové RJ, Medeiros TX, Annex BH, Popel AS. A mechanistic integrative computational model of macrophage polarization: Implications in human pathophysiology. PLoS Comput Biol. 2019;15: 1–28. doi:10.1371/journal.pcbi.1007468

17. Raser JM, O’Shea EK. Control of Stochasticity in Eukaryotic Gene Expression. Science (80-). 2004;304: 1811–1814. doi:10.1126/science.1098641

18. Dey SS, Foley JE, Limsirichai P, Schaffer D V., Arkin AP. Orthogonal control of expression mean and variance by epigenetic features at different genomic loci. Mol Syst Biol. 2015;11: 806. doi:10.15252/msb.20145704

19. Bartman CR, Hamagami N, Keller CA, Giardine B, Hardison RC, Blobel GA, et al. Transcriptional Burst Initiation and Polymerase Pause Release Are Key Control Points of Transcriptional Regulation. Mol Cell. 2019;73: 519–532.e4. doi:10.1016/J.MOLCEL.2018.11.004

20. Peccoud J, Ycart B. Markovian modeling of gene-product synthesis. Theor Popul Biol. 1995;48: 222–234. doi:10.1006/tpbi.1995.1027

21. Bahar Halpern K, Tanami S, Landen S, Chapal M, Szlak L, Hutzler A, et al. Bursty gene expression in the intact mammalian liver. Mol Cell. 2015;58: 147–156. doi:10.1016/j.molcel.2015.01.027

22. Tantale K, Mueller F, Kozulic-Pirher A, Lesne A, Victor J-M, Robert M-C, et al. A single-molecule view of transcription reveals convoys of RNA polymerases and multi-scale bursting. Nat Commun. 2016;7: 12248. doi:10.1038/ncomms12248

23. Rodriguez J, Ren G, Day CR, Zhao K, Chow CC, Larson DR. Intrinsic Dynamics of a Human Gene Reveal the Basis of Expression Heterogeneity. Cell. 2019;176: 213–226.e18. doi:10.1016/j.cell.2018.11.026

24. Corrigan AM, Tunnacliffe E, Cannon D, Chubb JR. A continuum model of transcriptional bursting. Elife. 2016;5: e13051. doi:10.7554/eLife.13051

25. Harper C V., Finkenstädt B, Woodcock DJ, Friedrichsen S, Semprini S, Ashall L, et al. Dynamic Analysis of Stochastic Transcription Cycles. Levchenko A, editor. PLoS Biol. 2011;9: e1000607. doi:10.1371/journal.pbio.1000607

26. Zoller B, Nicolas D, Molina N, Naef F. Structure of silent transcription intervals and noise characteristics of mammalian genes. Mol Syst Biol. 2015;11: 823. doi:10.15252/msb.20156257

27. Friedrich D, Friedel L, Finzel A, Herrmann A, Preibisch S, Loewer A. Stochastic transcription in the p53-mediated response to &lt;scp>DNA&lt;/scp> damage is modulated by burst frequency. Mol Syst Biol. 2019;15: 1–20. doi:10.15252/msb.20199068

28. Bass VL, Wong VC, Bullock ME, Gaudet S, Miller-Jensen K. TNF stimulation primarily modulates transcriptional burst size of NF-κB-regulated genes. Mol Syst Biol. 2021;17: e10127. doi:10.15252/msb.202010127

29. Pahl HL. Activators and target genes of Rel/NF-κB transcription factors. Oncogene. 1999;18: 6853–6866. doi:10.1038/sj.onc.1203239

30. Zhong H, May MJ, Jimi E, Ghosh S. The phosphorylation status of nuclear NF-κB determines its association with CBP/p300 or HDAC-1. Mol Cell. 2002;9: 625–636. doi:10.1016/S1097-2765(02)00477-X

31. van Essen D, Engist B, Natoli G, Saccani S. Two Modes of Transcriptional Activation at Native Promoters by NF-κB p65. Ashwell JD, editor. PLoS Biol. 2009;7: e1000073. doi:10.1371/journal.pbio.1000073

32. Yang Z, Yik JHN, Chen R, He N, Moon KJ, Ozato K, et al. Recruitment of P-TEFb for stimulation of transcriptional elongation by the bromodomain protein Brd4. Mol Cell. 2005;19: 535–545. doi:10.1016/j.molcel.2005.06.029

33. Itzen F, Greifenberg AK, Bösken CA, Geyer M. Brd4 activates P-TEFb for RNA polymerase II CTD phosphorylation. Nucleic Acids Res. 2014;42: 7577–7590. doi:10.1093/nar/gku449

34. Brasier AR. Perspective: Expanding role of cyclin dependent kinases in cytokine inducible gene expression. Cell Cycle. 2008;7: 2661–2666. doi:10.4161/cc.7.17.6594

35. Rogatsky I, Adelman K. Preparing the first responders: building the inflammatory transcriptome from the ground up. Mol Cell. 2014;54: 245–54. doi:10.1016/j.molcel.2014.03.038

36. Barboric M, Nissen RMRM, Kanazawa S, Jabrane-Ferrat N, Peterlin BM. NF-κB Binds P-TEFb to Stimulate Transcriptional Elongation by RNA Polymerase II. Mol Cell. 2001;8: 327–337. doi:10.1016/S1097-2765(01)00314-8

37. Weinberger L, Voichek Y, Tirosh I, Hornung G, Amit I, Barkai N. Expression Noise and Acetylation Profiles Distinguish HDAC Functions. Mol Cell. 2012;47: 193–202. doi:10.1016/j.molcel.2012.05.008

38. Burnett JC, Miller-Jensen K, Shah PS, Arkin AP, Schaffer D V. Control of Stochastic Gene Expression by Host Factors at the HIV Promoter. Bieniasz PD, editor. PLoS Pathog. 2009;5: e1000260. doi:10.1371/journal.ppat.1000260

39. Gressel S, Schwalb B, Decker TM, Qin W, Leonhardt H, Eick D, et al. CDK9-dependent RNA polymerase II pausing controls transcription initiation. Elife. 2017;6. doi:10.7554/eLife.29736

40. Shao W, Zeitlinger J. Paused RNA polymerase II inhibits new transcriptional initiation. Nat Genet. 2017;49: 1045–1051. doi:10.1038/ng.3867

41. Craigie R, Bushman FD. HIV DNA integration. Cold Spring Harb Perspect Med. 2012;2: a006890. doi:10.1101/cshperspect.a006890

42. Finzi D, Hermankova M, Pierson T, Carruth LM, Buck C, Chaisson RE, et al. Identification of a reservoir for HIV-1 in patients on highly active antiretroviral therapy. Science (80-). 1997;278: 1295–1300. doi:10.1126/science.278.5341.1295

43. Chan JKL, Greene WC. NF-κB/Rel: agonist and antagonist roles in HIV-1 latency. Curr Opin HIV AIDS. 2011;6: 12–18. doi:10.1097/COH.0b013e32834124fd

44. Hayden MS, Ghosh S. Shared Principles in NF-κB Signaling. Cell. 2008;132: 344–362. doi:10.1016/j.cell.2008.01.020

45. Skupsky R, Burnett JC, Foley JE, Schaffer D V., Arkin AP. HIV Promoter Integration Site Primarily Modulates Transcriptional Burst Size Rather Than Frequency. Friedman N, editor. PLoS Comput Biol. 2010;6: e1000952. doi:10.1371/journal.pcbi.1000952

46. Jordan A, Defechereux P, Verdin E. The site of HIV-1 integration in the human genome determines basal transcriptional activity and response to Tat transactivation. EMBO J. 2001;20: 1726–1738. doi:10.1093/emboj/20.7.1726

47. Miller-Jensen K, Dey SS, Schaffer D V., Arkin AP. Varying virulence: epigenetic control of expression noise and disease processes. Trends Biotechnol. 2011;29: 517–525. doi:10.1016/j.tibtech.2011.05.004

48. Miller-Jensen K, Skupsky R, Shah PS, Arkin AP, Schaffer D V. Genetic Selection for Context-Dependent Stochastic Phenotypes: Sp1 and TATA Mutations Increase Phenotypic Noise in HIV-1 Gene Expression. Covert MW, editor. PLoS Comput Biol. 2013;9: e1003135. doi:10.1371/journal.pcbi.1003135

49. Chavali AK, Wong VC, Miller-Jensen K. Distinct promoter activation mechanisms modulate noise-driven HIV gene expression. Sci Rep. 2015;5: 17661. doi:10.1038/srep17661

50. Corrigan AM, Chubb JR. Regulation of Transcriptional Bursting by a Naturally Oscillating Signal. Curr Biol. 2014;24: 205–211. doi:10.1016/j.cub.2013.12.011

51. Raj A, van Oudenaarden A. Nature, Nurture, or Chance: Stochastic Gene Expression and Its Consequences. Cell. 2008;135: 216–226. doi:10.1016/j.cell.2008.09.050

52. Larsson AJM, Johnsson P, Hagemann-Jensen M, Hartmanis L, Faridani OR, Reinius B, et al. Genomic encoding of transcriptional burst kinetics. Nature. Nature Publishing Group; 2019. pp. 251–254. doi:10.1038/s41586-018-0836-1

53. Kwak H, Fuda NJ, Core LJ, Lis JT. Precise Maps of RNA Polymerase Reveal How Promoters Direct Initiation and Pausing. Science (80-). 2013;339: 950–953. doi:10.1126/science.1229386

54. Rahl PB, Lin CY, Seila AC, Flynn RA, McCuine S, Burge CB, et al. c-Myc Regulates Transcriptional Pause Release. Cell. 2010;141: 432–445. doi:10.1016/j.cell.2010.03.030

55. Muse GW, Gilchrist DA, Nechaev S, Shah R, Parker JS, Grissom SF, et al. RNA polymerase is poised for activation across the genome. Nat Genet. 2007;39: 1507–1511. doi:10.1038/ng.2007.21

56. Jonkers I, Kwak H, Lis JT. Genome-wide dynamics of Pol II elongation and its interplay with promoter proximal pausing, chromatin, and exons. Elife. 2014;3. doi:10.7554/eLife.02407

57. Wakamori M, Okabe K, Ura K, Funatsu T, Takinoue M, Umehara T. Quantification of the effect of site-specific histone acetylation on chromatin transcription rate. Nucleic Acids Res. 2020;48: 12648–12659. doi:10.1093/nar/gkaa1050

58. Bryan LC, Weilandt DR, Bachmann AL, Kilic S, Lechner CC, Odermatt PD, et al. Single-molecule kinetic analysis of HP1-chromatin binding reveals a dynamic network of histone modification and DNA interactions. Nucleic Acids Res. 2017;45: 10504–10517. doi:10.1093/nar/gkx697

59. Lenstra TL, Rodriguez J, Chen H, Larson DR. Transcription Dynamics in Living Cells. Annu Rev Biophys. 2016;45: 25–47. doi:10.1146/annurev-biophys-062215-010838

60. Cao Y, Lei X, Ribeiro RM, Perelson AS, Liang J. Probabilistic control of HIV latency and transactivation by the Tat gene circuit. Proc Natl Acad Sci. 2018; 201811195. doi:10.1073/pnas.1811195115

61. Wang Q, Lin J. Heterogeneous recruitment abilities to RNA polymerases generate nonlinear scaling of gene expression with cell volume. Nat Commun. 2021;12: 6852. doi:10.1038/s41467-021-26952-y

62. Rosen GA, Baek I, Friedman LJ, Joo YJ, Buratowski S, Gelles J. Dynamics of RNA polymerase II and elongation factor Spt4/5 recruitment during activator-dependent transcription. Proc Natl Acad Sci. 2020;117: 32348–32357. doi:10.1073/pnas.2011224117

63. Zobeck KL, Buckley MS, Zipfel WR, Lis JT. Recruitment Timing and Dynamics of Transcription Factors at the Hsp70 Loci in Living Cells. Mol Cell. 2010;40: 965–975. doi:10.1016/j.molcel.2010.11.022

64. Palangat M, Meier TI, Keene RG, Landick R. Transcriptional Pausing at +62 of the HIV-1 Nascent RNA Modulates Formation of the TAR RNA Structure. Mol Cell. 1998;1: 1033–1042. doi:10.1016/S1097-2765(00)80103-3

65. He N, Liu M, Hsu J, Xue Y, Chou S, Burlingame A, et al. HIV-1 Tat and host AFF4 recruit two transcription elongation factors into a bifunctional complex for coordinated activation of HIV-1 transcription. Mol Cell. 2010;38: 428–38. doi:10.1016/j.molcel.2010.04.013

66. Sobhian B, Laguette N, Yatim A, Nakamura M, Levy Y, Kiernan R, et al. HIV-1 Tat Assembles a Multifunctional Transcription Elongation Complex and Stably Associates with the 7SK snRNP. Mol Cell. 2010;38: 439–451. doi:10.1016/j.molcel.2010.04.012

67. Nabel G, Baltimore D. An inducible transcription factor activates expression of human immunodeficiency virus in T cells. Nature. 1987;326: 711–713. doi:10.1038/326711a0

68. Weinberger LS, Burnett JC, Toettcher JE, Arkin AP, Schaffer D V. Stochastic gene expression in a lentiviral positive-feedback loop: HIV-1 Tat fluctuations drive phenotypic diversity. Cell. 2005;122: 169–82. doi:10.1016/j.cell.2005.06.006

69. Miller-Jensen K, Dey SS, Pham N, Foley JE, Arkin AP, Schaffer D V. Chromatin accessibility at the HIV LTR promoter sets a threshold for NF-κB mediated viral gene expression. Integr Biol. 2012;4: 661. doi:10.1039/c2ib20009k

70. Razooky BS, Cao Y, Hansen MMK, Perelson AS, Simpson ML, Weinberger LS. Nonlatching positive feedback enables robust bimodality by decoupling expression noise from the mean. PLoS Biol. 2017;15: 1–26. doi:10.1371/journal.pbio.2000841

71. Fraser LTCR, Dikdan RJ, Dey S, Singh A, Tyagi S. Reduction in gene expression noise by targeted increase in accessibility at gene loci. Proc Natl Acad Sci U S A. 2021;118. doi:10.1073/pnas.2018640118

72. Ma L, Gao Z, Wu J, Zhong B, Xie Y, Huang W, et al. Co-condensation between transcription factor and coactivator p300 modulates transcriptional bursting kinetics. Mol Cell. 2021;81: 1682–1697.e7. doi:10.1016/j.molcel.2021.01.031

73. Wong VC, Mathew S, Ramji R, Gaudet S, Miller-Jensen K. Fold-Change Detection of NF-κB at Target Genes with Different Transcript Outputs. Biophys J. 2019;116: 709–724. doi:10.1016/j.bpj.2019.01.011

74. Kumar N, Singh A, Kulkarni R V. Transcriptional Bursting in Gene Expression: Analytical Results for General Stochastic Models. Morozov A V, editor. PLOS Comput Biol. 2015;11: e1004292. doi:10.1371/journal.pcbi.1004292

75. Reuse S, Calao M, Kabeya K, Guiguen A, Gatot J-S, Quivy V, et al. Synergistic Activation of HIV-1 Expression by Deacetylase Inhibitors and Prostratin: Implications for Treatment of Latent Infection. Sommer P, editor. PLoS One. 2009;4: e6093. doi:10.1371/journal.pone.0006093

76. Wong VC, Fong LE, Adams NM, Xue Q, Dey SS, Miller-Jensen K. Quantitative evaluation and optimization of co-drugging to improve anti-HIV latency therapy. Cell Mol Bioeng. 2014;7: 320–333. doi:10.1007/s12195-014-0336-9

77. Sneddon MW, Faeder JR, Emonet T. Efficient modeling, simulation and coarse-graining of biological complexity with NFsim. Nat Methods. 2011;8: 177–183. doi:10.1038/nmeth.1546

78. Gillespie DT. Exact stochastic simulation of coupled chemical reactions. J Phys Chem. 1977;81: 2340–2361. doi:10.1021/j100540a008

79. Hartigan PM. Algorithm AS 217: Computation of the Dip Statistic to Test for Unimodality. Appl Stat. 1985;34: 320. doi:10.2307/2347485

80. Hartigan JA, Hartigan PM. The Dip Test of Unimodality. Ann Stat. 1985;13: 70–84. doi:10.1214/aos/1176346577

81. Harris LA, Hogg JS, Tapia J-J, Sekar JAP, Gupta S, Korsunsky I, et al. BioNetGen 2.2: advances in rule-based modeling. Bioinformatics. 2016;32: 3366–3368. doi:10.1093/bioinformatics/btw469

